# A partner-resolved marine sponge hologenome reveals that three bacterial symbionts disproportionately expand holobiont metabolism via enriched membrane transporter repertoires

**DOI:** 10.64898/2026.08.05.743132

**Authors:** Xueyan Xiang, Eve Maunders, Bernard M Degnan, Sandie M Degnan

**Affiliations:** Centre for Marine Science and School of the Environment, The University of Queensland, Brisbane QLD 4072, Australia

**Keywords:** *Amphimedon queenslandica*, hologenomics, holotranscriptomics, host-microbe interactions, marine symbionts, metabolome, sponge microbiome

## Abstract

Marine sponges associate with microbial symbionts that shape host physiology and drive nutrient cycling, yet assigning partner-specific contributions to holobiont metabolism remains challenging in species with complex microbiomes. The coral reef demosponge *Amphimedon queenslandica* offers tractable resolution, as its adult microbiome is dominated by just three extracellular, vertically inherited gammaproteobacterial symbionts (*AqS1*, *AqS2* and *AqS3*). Here, we present improve genome assemblies of the host and each symbiont, integrated with adult holotranscriptomes to resolve functional partitioning and expressed pathway capacity of each partner. Despite representing only ∼20% of KEGG-annotated hologenome genes, the three symbionts contribute 45.2 and 72% of metabolic and membrane transporter genes, respectively, revealing a pronounced functional imbalance in proteins involved in nutrient transformation and exchange. Although central heterotrophic carbon metabolism is complete across all four partners, gene content and expression are consistent with symbiont uptake of host-liberated carbohydrates. Complementation of pathways occurs across dissolved inorganic nutrient assimilation, including nitrate reduction, sulfur redox metabolism, and phosphate uptake and storage. Symbionts further expand holobiont biosynthetic breadth through amino acid, vitamin, and co-factor pathways that are incomplete or absent from the host, coupled with expressed membrane transporter repertoires consistent with directed metabolite exchange. Together, these results show that vertically inherited symbionts can disproportionately expand holobiont metabolic capacity, with membrane transporter enrichment linking symbiont metabolic breadth to host physiology. This exchange-oriented functional architecture, rather than simple pathway redundancy, appears to underlie metabolic integration in this low-complexity animal– microbe symbiosis.

## Introduction

Marine sponges (phylum Porifera) are functionally important components of benthic ecosystems from the tropics to the poles (Bell 2008). Through high biomass and a sessile, filter-feeding lifestyle, sponges contribute to benthic–pelagic coupling, bioerosion and substrate stabilisation, with particularly strong ecosystem impacts in oligotrophic habitats such as coral reefs (de Goeij et al. 2013; Folkers and Rombouts 2020). By filtering thousands of litres of seawater per kilogram of biomass per day (Bell 2008; Leys et al. 2011), sponges capture both particulate food and dissolved organic matter, the latter of which is the most abundant heterotrophic resource in the ocean (de Goeij et al. 2013; Rix et al. 2016, 2017, 2020). A substantial fraction of assimilated material is subsequently released as particulate organic matter via cell shedding, making sponge-processed nutrients available to higher trophic levels and contributing to reef-wide nutrient retention and recycling (de Goeij et al. 2013; Maldonado 2015; Pawlik et al. 2016).

A defining feature of many sponges is their association with often dense and phylogenetically diverse microbial communities that extend host physiology and mediate major biogeochemical transformations (Webster and Thomas 2016; O’Brien et al. 2020; Steinert et al. 2020). Sponge-associated microbes encode broad metabolic capabilities, including heterotrophic carbon metabolism and, in some systems, autotrophic carbon fixation (Kamke et al. 2013; Li et al. 2015; Gauthier et al. 2016; Moitinho-Silva et al. 2017; Engelberts et al. 2020), as well as pathways for nitrogen fixation, ammonia oxidation, nitrate reduction and denitrification, influencing transformation and retention of dissolved inorganic and organic nitrogen (Fiore et al. 2013; Moitinho-Silva et al. 2017; Zhang et al. 2019). Beyond elemental cycling, symbionts may complement host metabolism through biosynthesis of amino acids, vitamins and cofactors that the host cannot produce de novo (Fiore et al. 2015; Gauthier et al. 2016; Bayer et al. 2018; Engelberts et al. 2020).

Despite this breadth of evidence, attributing specific biosynthetic functions to individual microbial partners remains a challenge. In many sponge systems, microbiome complexity combined with the absence of a well-annotated host genome makes it difficult to distinguish genuine metabolic complementation from pathway redundancy, and to determine which organisms encode and express factors that function in exchange of substrates and products between partners. Membrane transport is particularly under interpreted in community-level inventories, yet it is central to metabolite trafficking and therefore to the question of whether symbionts actively provision the host or simply co-occur metabolically. Holobionts that combine low microbial complexity, partner-resolved genomes, and matched expression data offer the most tractable route to answering this question.

The coral reef demosponge *Amphimedon queenslandica* provides such a system. Its microbiome is dominated by three extracellular, heterotrophic bacterial symbionts (hereafter *AqS1*, *AqS2* and *AqS3*) that are maternally-inherited, persist stably across seasons and life cycle stages, and together comprise 90% or more of adult bacterial abundance (Fieth et al. 2016; Gauthier et al. 2016). *AqS1* is a sulfur-oxidizing gammaproteobacterium in the order Chromatiales that can alone exceed 60% of the adult microbiome (Fieth et al. 2016; Gauthier et al. 2016; Yang et al. 2026). *AqS2* (*Amphirhobacter heronislandensis*) belongs to the recently described gammaproteobacterial order Tethybacterales (Taylor et al. 2021), while *AqS3* represents another divergent gammaproteobacterial lineage (Gauthier et al. 2016). Draft genomes are available for all three symbionts (Gauthier et al. 2016), and the *A. queenslandica* host genome is extensively annotated (Srivastava et al. 2010; Fernandez-Valverde et al. 2015), enabling direct comparisons of gene content and expression between holobiont partners.

Here, we improve genome assemblies and annotations for *A. queenslandica* and *AqS1*–*AqS3*, and integrate these with adult holobiont transcriptomes to resolve metabolic capacities and functional partitioning at partner-level resolution. We show that three vertically inherited symbionts disproportionately expand holobiont metabolic capacity, not through numerical dominance of gene content, but through enrichment of those functional categories most relevant to nutrient transformation and exchange, namely metabolism and membrane transport. This transport-enriched, exchange-oriented functional architecture appears to be the mechanistic basis of metabolic integration in this low-complexity animal– microbe symbiosis.

## Materials and methods

### Hologenome sequencing, assembly and annotation

Genomic DNA was extracted from two adult *Amphimedon queenslandica* collected from Heron Island Reef, Australia (23.44°S, 151.92°E) under GBRMPA permit G16/38120.1 as described previously (Srivastava et al. 2010). For one individual, three in vitro chromatin reconstituted “Chicago” libraries (2 × 101 bp; insert sizes 1–300 kb) were prepared and sequenced on an Illumina HiSeq2000 platform by Dovetail Genomics (USA) (Putnam et al. 2016). For the other individual, a TruSeq Nano DNA LT library (insert size ∼350 bp) was sequenced on an Illumina NextSeq500 platform across four lanes (2 × 75 bp paired-end reads).

Genome reassembly of *A. queenslandica* and the three dominant symbionts was performed using the HiRise scaffolding pipeline (Putnam et al. 2016). Chicago and Illumina reads were aligned to the existing host (Srivastava et al. 2010; Fernandez-Valverde et al. 2015) and symbiont reference genomes (Gauthier et al. 2016) using a modified version of SNAP (Putnam et al. 2016). Species-specific read pairs were extracted with BEDTools v2.26.0 (Quinlan and Hall 2010) prior to scaffolding with HiRise (version HiRise_July2015_GR). Assembly completeness was assessed using BUSCO v3.0.2 (Simão et al. 2015) with the metazoa_odb9 lineage dataset for the host and proteobacteria_odb9 for symbionts.

Host gene models were generated by liftover of Aqu2.1 annotations (Fernandez-Valverde et al. 2015) onto the new assembly using FLO (Pracana et al. 2017), yielding the Aqu3.1 gene model set. Symbiont genomes were annotated de novo using Prokka v1.12 (Seemann 2014) with default parameters, yielding *AqS1*v2, *AqS2*v2, and *AqS3*v2 gene model sets. Functional annotation of all four partner genomes was performed using Blast2GO v5.2.4 (Conesa and Götz 2008), with Blastp-fast searches against the NCBI non-redundant (nr) protein database (e-value threshold ≤ 1×10⁻³). The improved host and symbiont genome assemblies and annotations have been deposited in NCBI [BioProject PRJNA668660].

### KEGG module reconstruction and identification of symbiont modules

KEGG Orthology (KO) assignments were obtained using GhostKOALA (Kanehisa et al. 2016) against the KEGG GENES database, using the family_eukaryotes gene set for the host and genus_prokaryotes for each symbiont. KofamKOALA (Aramaki et al. 2020) and DeepKOALA (Yu et al. 2026) were also performed for the KO annotation with default parameters. The final KO assignments were the combined results of the four functional annotation methods and used for all subsequent KEGG pathway and module analyses.

KEGG module completeness for the holobiont and each species was calculated with the kegg-pathways-completeness tool from MGnify (Burgin et al. 2023). The modules with 100% completeness in the holobiont but incomplete in the host alone were identified as candidate symbiont-supported modules. Candidate modules were retained if there was implied functional exchange or complementation between host and symbiont rather than purely prokaryote-internal metabolism with no plausible connection to host physiology. Each candidate module was manually inspected to confirm step-level completeness within individual symbiont genomes and to verify that key enzymatic functions were not misassigned across phylogenetically distant homologues.

### Holotranscriptome sequencing and analysis

Six *A. queenslandica* adults were collected from Heron Island Reef and maintained in flow-through aquaria under ambient water, temperature and light conditions for no more than 12 hours prior to sampling. Tissue biopsies (∼3 cm³) were taken from each individual and processed to enrich for bacterial cells by sequential size-selective filtration followed by low-speed centrifugation, as described in Thomas et al. (2010). Total RNA was extracted using TRIzol (Thermo Fisher Scientific) following the manufacturer’s protocol. RNA quantity was assessed by Qubit fluorometric quantification and RNA integrity was confirmed by Agilent Bioanalyzer.

Strand-specific libraries were prepared using an Illumina TruSeq Stranded Total RNA protocol modified as follows. To enable recovery of both host and symbiont transcripts, we replaced the more usual poly(A) selection with a simultaneous depletion of both sponge and bacterial symbiont ribosomal RNA conducted using a custom-designed 3’-biotinylated DNA oligos riboPOOL kit (siTOOLs Biotech) following the manufacturer’s protocol. Details of our kit are available at https://www.sitoolsbiotech.com/products/ribopools/rrna-depletion/available-ribopools/amphimedon-queenslandica. Strand specificity was retained via dUTP incorporation. Libraries were sequenced on an Illumina NovaSeq 6000 to generate paired-end reads. These six transcriptomes were combined with three previously published rRNA-depleted holobiont RNA-Seq datasets (Xiang et al. 2022), yielding a combined dataset of nine adult *A. queenslandica* holotranscriptomes for all downstream analyses.

Raw reads from the nine holotranscriptomes were quality-filtered and adapter-trimmed using Trimmomatic v0.36 (Bolger et al. 2014) with standard parameters. Trimmed reads were aligned simultaneously to all four partner genomes (Aqu3.1, AqS1v2, AqS2v2, and AqS3v2) using HISAT2 v2.0.5 (Kim et al. 2015). Reads mapping to more than one genome were discarded to ensure species-specific attribution. Species-specific alignment files were generated using SAMtools v1.3 (Li et al. 2009). Gene-level read counts were obtained using htseq-count v0.11.2 (Anders et al. 2015), with strandedness set to "reverse". Genes with at least one count in at least three of the nine samples were retained. Gene expression levels were normalized to transcripts per million (TPM) (Wagner et al. 2012). Expressed genes within each partner were ranked by mean TPM across the nine holotranscriptomes and divided into equal expression quartiles, with quartile 1 (Q1) containing the lowest expressed genes and Q4 the highest.

Pathway-level transcriptional activity was assessed using expressed KO-annotated genes mapped onto KEGG modules and pathways. For symbionts, a module was considered transcriptionally active if more than 60% of module component genes were expressed and all enzymatic steps designated as key reactions by KEGG were expressed, following the criteria of Engelberts et al. (2020). Where multiple genes encoded the same KO function within a single genome, the copy with the highest mean TPM was used as the representative for pathway visualisation and module activity assessment.

## Results

### Improved hologenome assemblies provide partner-resolved functional resolution

To resolve how individual symbionts contribute to holobiont metabolism, we first improved genome assemblies for *Amphimedon queenslandica* and its three dominant bacterial symbionts (*AqS1*, *AqS2* and *AqS3*) by integrating long-range chromatin data with short-read Illumina sequencing (Table 1 and Tables S1-S3). These new assemblies, combined with the low complexity of the *A. queenslandica* microbiome, improved the assignment of genes, pathways, and transcript profiles to individual holobiont partners. Improved contiguity yielded more complete coding sequences (CDS), particularly for the symbiont genomes, with *AqS1* having fewer but longer CDS, and *AqS2* and *AqS3* having more CDS (Tables S3 and S4). The new *A. queenslandica* assembly (Aqu3.1) also yielded fewer CDS, of which 86.4% were functionally annotated by Blast2GO (Table 1 and Tables S3 and S5). BUSCO completeness scores were maintained or improved across all assemblies. The three symbiont genomes together contribute 15.8% of total hologenome CDS.

**Table 1.** Genome assembly statistics of *A. queenslandica* (*Aqu*), and three bacterial symbionts, *AqS1*, *AqS2* and *AqS3*.

| Genome version | *Aqu* |  | *AqS1* |  | *AqS2* |  | *AqS3* |  |
| --- | --- | --- | --- | --- | --- | --- | --- | --- |
|  | Aqu2.1 | Aqu3.1 | v1 | v2 | v1 | v2 | v1 | v2 |
| Total length (Mb) | 166.7 | 167.7 | 4.20 | 4.21 | 1.61 | 1.63 | 3.16 | 3.17 |
| Scaffold number | 13,397 | 3,871 | 127 | 116 | 239 | 68 | 233 | 151 |
| Max. scaffold length (kb) | 1,889 | 4,599 | 266 | 288 | 83 | 441 | 173 | 312 |
| Scaffold N50 length (kb) | 120.4 | 950.5 | 79.3 | 102.7 | 12.2 | 147.7 | 35.2 | 90.0 |
| % BUSCO completeness | 86.8 | 86.6 | 95.0 | 95.1 | 64.7 | 69.2 | 89.1 | 89.6 |
| CDS number | 44,001 | 42,644 | 3,767 | 3,478 | 1,349 | 1,621 | 2,418 | 2,933 |
| % hologenome CDS | 85.4 | 84.2 | 7.3 | 6.9 | 2.6 | 3.2 | 4.7 | 5.8 |

The Aqu2.1 and v1 assemblies are previously published *A. queenslandica* and symbiont annotated genome assemblies, respectively (Srivastava et al. 2010; Fernandez-Valverde et al. 2015; Gauthier et al. 2016), and Aqu3.1 and v2 are the improved genome assemblies (this study).

### Symbionts disproportionately contribute to holobiont metabolism and membrane transport

Functional annotation using KEGG revealed that although *AqS1*, *AqS2*, and *AqS3* together account for only 20.4% of KEGG-annotated hologenome genes, they encode 43.9% of KEGG-annotated metabolic genes (Fig. 1, and Tables S6 and S7). This enrichment is not confined to a narrow functional niche, but it spans nearly all KEGG metabolic subcategories (Fig. 1B). In addition, the symbionts encode 72% of holobiont membrane transporter genes (Fig. 1B). *AqS1* and *AqS2* together account for the majority of these, with both encoding large repertoires spanning sugar uptake, amino acid transport, inorganic ion handling, and vitamin import and export systems. *AqS3* contributes a smaller but complementary transporter load consistent with its more restricted metabolic role. The host, by contrast, encodes 28% of holobiont membrane transporter genes despite encoding approximately 79.6% of total KEGG-annotated hologenome genes, a disparity that reflects the concentration of exchange-enabling capacity in symbiont genomes and underpins the exchange-oriented architecture described in the sections below. This disproportionate contribution of metabolic and transport genes in symbionts is consistent with them not merely being metabolically capable but being structurally positioned to acquire substrates and products, redistribute metabolic intermediates, and provision products to the host.

**Fig. 1.**
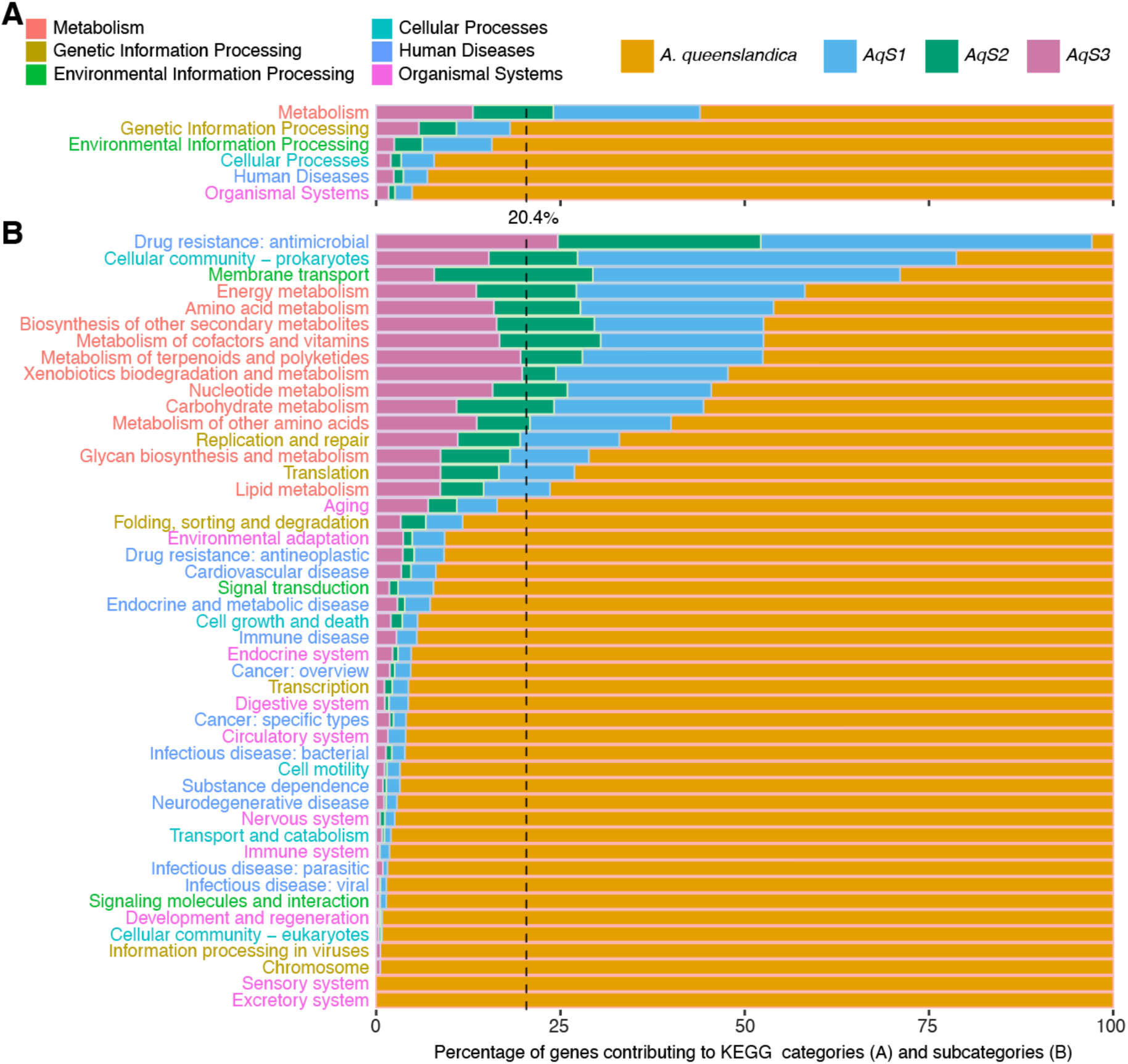
Functional partitioning in the *A. queenslandica* hologenome. **(A)** Percent genes contributed by *A. queenslandica*, and symbionts *AqS1*, *AqS2* and *AqS3* to broad KEGG biological categories. **(B)** Percent genes contributed to KEGG subcategories. Y-label colours of sub-categories in B are consistent with y-label colours of categories in A. Of the total hologenome genes that could be assigned to a KO group, 20.4% are from the three bacterial symbiont genomes (dashed line; Table S6).

In contrast, signalling pathways and other multicellular systems are largely restricted to the sponge host. This is consistent with a broad division of labour in which symbionts appear to specialise in metabolic breadth and exchange while the host appears to retain regulatory and physiological control (Fig. 1). Antimicrobial resistance genes were encoded exclusively by symbionts, with contributions from all three genomes (Fig. 1B).

Nine adult holobiont transcriptomes confirmed that these genome-predicted capacities are active, with 73.8% of *A. queenslandica* genes and 92.1, 95.9 and 81.6% of *AqS1*, *AqS2*, and *AqS3*, respectively, being expressed (Table S8). The proportion of RNA-seq reads mapping to symbiont genomes reflects symbiont abundance in the adult microbiome, with *AqS1*, *AqS2*, and *AqS3* accounting for 19.01, 1.25 and 0.87% of uniquely mapping reads, respectively (Table S9). Genes underpinning symbiont-enriched functional categories, including metabolism and membrane transport, were consistently expressed at moderate to high levels (Quartiles 2-4; Q2–Q4) in all transcriptomes, confirming that the functional imbalances observed at the genome level are transcriptionally deployed in adults (Tables S10 and S11).

### Pathway-level evidence for transport-enabled division of labour

KEGG reconstruction identified 37 modules across 25 pathways that are incomplete in *A. queenslandica* alone but complete at the holobiont level through symbiont contributions. These symbiont-supported modules were enriched for amino acid metabolism, cofactor and vitamin metabolism, glycan biosynthesis and metabolism, carbohydrate metabolism, lipid metabolism, and energy metabolism (Table 2 and Table S12). In the following sections we examine representative pathways across carbon metabolism, amino acid biosynthesis, vitamin and cofactor production, and inorganic nutrient assimilation, focussing on how symbiont metabolic capacity and membrane transporter repertoires can connect symbiont metabolic breadth to host physiology. In each case, pathway-level expression data confirm that symbiont contributions are actively deployed in the adult sponge.

**Table 2.** KEGG pathways that are incomplete in *A. queenslandica* alone, but complete in the holobiont.

| Functional category | Modules | Pathways |
| --- | --- | --- |
| Amino acid metabolism | 14 | Arg biosynthesis; Gly, Ser and Thr metabolism; Cys and Met metabolism; Val, Leu and Iso biosynthesis; Lys biosynthesis; His metabolism; Phe, Tyr and Trp biosynthesis |
| Cofactor and vitamin metabolism | 8 | Thiamine (vitamin B1) metabolism; Riboflavin (B2) metabolism; One carbon pool by folate; Nicotinate and nicotinamide metabolism; Pantothenate (B5) and CoA biosynthesis; Folate biosynthesis; Porphyrin metabolism |
| Glycan biosynthesis and metabolism | 5 | Amino sugar and nucleotide sugar metabolism; Biosynthesis of various nucleotide sugars |
| Carbohydrate metabolism | 5 | Citrate cycle (TCA cycle); Pentose phosphate pathway; Galactose metabolism; Glyoxylate and dicarboxylate metabolism; Propanoate metabolism |
| Lipid metabolism | 3 | Fatty acid biosynthesis; Glycerophospholipid metabolism |
| Energy metabolism | 2 | Oxidative phosphorylation; Sulfur metabolism |

The full list of 37 incomplete modules is provided in Table S12.

### Shared heterotrophic carbon metabolic core with symbiont-biased carbohydrate import

Genomic and transcriptional evidence indicates that the host and all three symbionts maintain aerobic heterotrophic metabolism. Glycolysis, pyruvate oxidation, the TCA (Krebs) cycle, and oxidative phosphorylation were complete or near-complete across the holobiont and broadly expressed in adults (Fig. 2, Tables S7, S10-S12 and Figs S1-5). Despite this shared metabolic core, partners differ in how they enter and sustain these pathways. These differences point toward a functional division between host substrate mobilisation and symbiont importation.

**Fig. 2.**
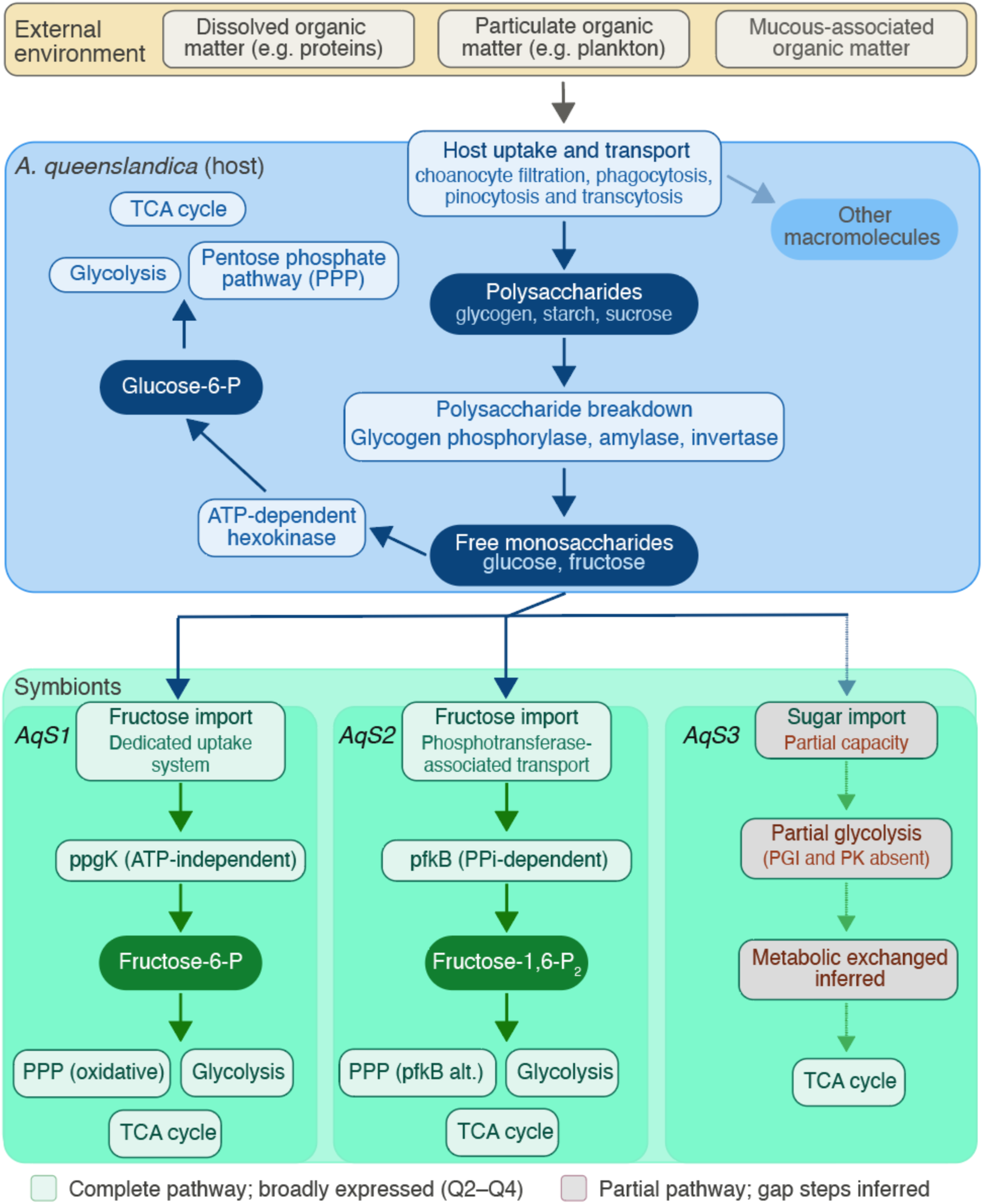
Carbon exchange in the *A. queenslandica* holobiont. Schematic summarising divergence points in carbon metabolism and inferred directional carbohydrate flow between *A. queenslandica* host and its symbionts (*AqS1*, *AqS2* and *AqS3*). Upper panel (tan) shows potential sources of external dietary carbon matter. Middle panel (blue) shows general sponge carbohydrate content, showing the consumption of external carbon via choanocyte filtration, phagocytosis and pinocytosis, and the mobilisation of stored polysaccharides by glycogen phosphorylase, amylase, and invertase to yield free monosaccharides. The host enters glycolysis via a canonical ATP-dependent hexokinase pathway. Lower panel (green) shows symbiont import and processing systems. *AqS1* encodes a dedicated fructose uptake system and phosphorylates imported fructose via ATP-independent polyphosphate glucokinase (ppgK). *AqS2* imports fructose via a phosphotransferase system and enters glycolysis via a pyrophosphate-dependent phosphofructokinase (pfkB). *AqS3* encodes partial sugar import capacity and is missing phosphoglucose isomerase (PGI) and pyruvate kinase (PK), suggesting partial reliance on metabolite exchange with other holobiont partners to sustain glycolytic flux. All four partners express complete or near-complete pyruvate oxidation and TCA cycle components (Figs S1-S3). The non-oxidative pentose phosphate pathway (PPP) branch is complete in the host and in *AqS1* (oxidative branch also complete in *AqS1*). *AqS2* and *AqS3* use an alternative route via pyrophosphate-dependent phosphofructokinase, compensating for the absence of transaldolase (Figs S4, S5). Solid arrows, inferred directional carbohydrate flow; smaller lighter arrows, pathway gaps imply dependence on intermediates from other partners; blue (host) and green (symbiont) blocks, complete and highly expressed (Q2-Q4) pathways; coral blocks, partial pathways; metabolites, dark coloured pills.

Entry into glycolysis diverges between partners, with *A. queenslandica* encoding a canonical hexokinase (EC 2.7.1.1) for glucose phosphorylation, and *AqS1* encoding polyphosphate glucokinase (EC 2.7.1.63, ppgK), which indicates an alternative phosphorylation route that does not consume ATP (Liao et al. 2012). *AqS3* shows additional absences, including phosphoglucose isomerase (EC 5.3.1.9) and pyruvate kinase (EC 2.7.1.40), suggesting partial reliance on alternative routes and/or metabolite exchange with other holobiont partners. The pentose phosphate pathway (PPP) was complete and strongly expressed in the host. In *AqS1* the oxidative branch was complete, while all three symbionts used pyrophosphate-dependent phosphofructokinase as an alternative non-oxidative PPP route, compensating for the absence of transaldolase (Tables S10 and S11).

The most functionally significant divergence between host and symbionts concerns carbohydrate mobilisation versus import. The host encodes and expresses enzymes for glycogen and starch mobilisation, and sucrose hydrolysis, consistent with liberation of simple sugars from stored or dietary polysaccharides. Symbionts, by contrast, express dedicated sugar import systems. *AqS1* encodes fructose uptake machinery and *AqS2* encodes phosphotransferase-associated fructose transport components (Fig. 2, Tables S10 and S12 and Figs S1, S2). This host-side liberation paired with symbiont-side import is consistent with directional carbohydrate flow from host to symbiont and illustrates the complementarity of the symbiont transporter repertoire and host metabolic activity.

### Symbiont amino acid biosynthesis and transport fills host gaps

The sponge host genome encodes incomplete biosynthetic pathways for 11 of the 20 standard amino acids (Table 2, Fig. 3, Tables S10-12 and Figs S6-13). Together, the three primary symbionts encode and express the pathways that are incomplete in the sponge, including for the biosynthesis of histidine, branched-chain amino acids (valine, leucine and isoleucine), and aromatic amino acids via the shikimate/chorismate pathway (phenylalanine, tryptophan and tyrosine). The host, in turn, encoded and expressed enzymes supporting synthesis of several non-essential amino acids, including asparagine, which was not supported by symbiont genomes.

**Fig. 3.**
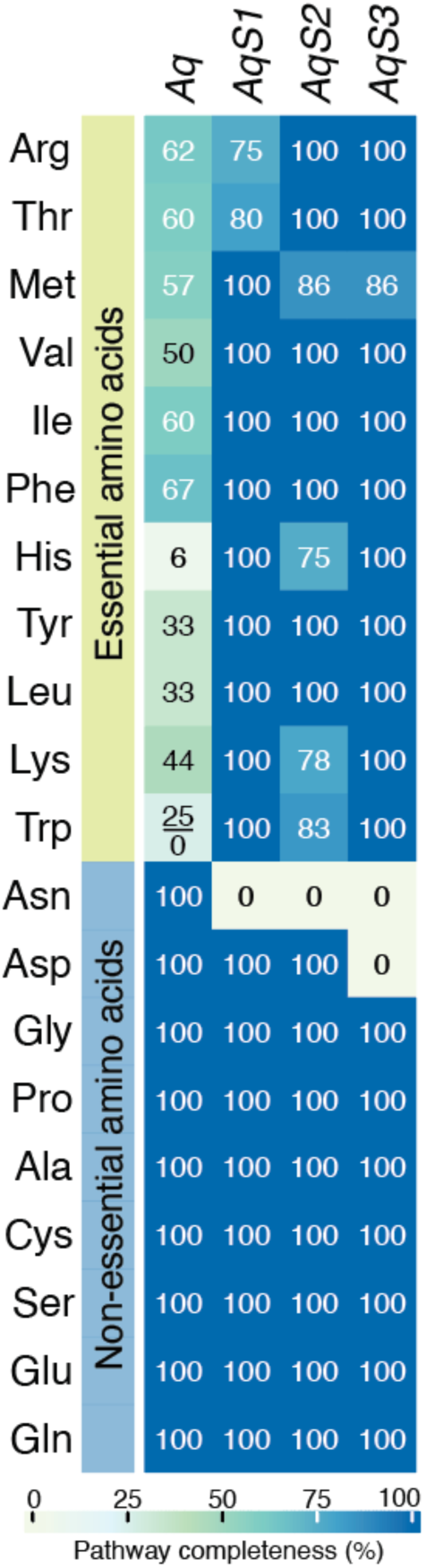
Amino acid biosynthesis in the *A. queenslandica* holobiont. Aq, *A. queenslandica*; *AqS1*, *AqS2* and *AqS3*, three primary symbionts. The heatmap and values show the percentage of enzymes present for each amino acid biosynthesis pathway encoded by each species and expressed in the adult (Figs S6-S13). The presence and expression of genes are the same in all cases except *A. queenslandica* tryptophan (Trp) genes, where the values above and below the line are percent genes in genome and expressed, respectively. Essential and nonessential amino acids in humans are annotated.

This complementary distribution of biosynthetic capacity is matched by a complementary distribution of transport functions. The host expressed multiple amino acid transporters, including L-type transporters consistent with uptake of a broad range of substrates. *AqS1* and *AqS2* encoded and expressed general and branched-chain amino acid transport systems (Tables S10-12). Together, these partner-resolved patterns — symbiont biosynthesis of amino acids the host cannot make, paired with expressed transporters on both sides — provide evidence for symbiont provisioning of essential amino acids to the host. These findings extend and contextualise prior isotope-tracing evidence for arginine complementation in *A. queenslandica* (Song et al. 2021) by placing it within a broader pattern of amino acid division of labour.

### Symbiont production of vitamins and cofactors is coupled with host uptake and within-pathway partitioning

Vitamin and cofactor pathways provided some of the strongest examples of division of labour, with symbiont biosynthetic capacity consistently paired with host transport and assimilation functions (Fig. 4, Tables S10-12 and Figs S14-16). Thiamine (vitamin B1) biosynthesis was supported primarily by *AqS1* and *AqS2*, while the host and *AqS3* encoded enzymes and transporters consistent with thiamine-phosphate conversion and/or uptake from the extracellular environment. Riboflavin (vitamin B2) biosynthesis was encoded by *AqS1* and *AqS3* and near-complete in *AqS2*, but entirely absent from *A. queenslandica*. The host instead encoded and expressed a riboflavin transporter, consistent with dependence on symbiont production and active uptake (Fig. 4 and Fig. S15). This pattern — symbiont biosynthesis, host transport — is among the clearest examples of obligate metabolic complementation in the holobiont.

**Fig. 4.**
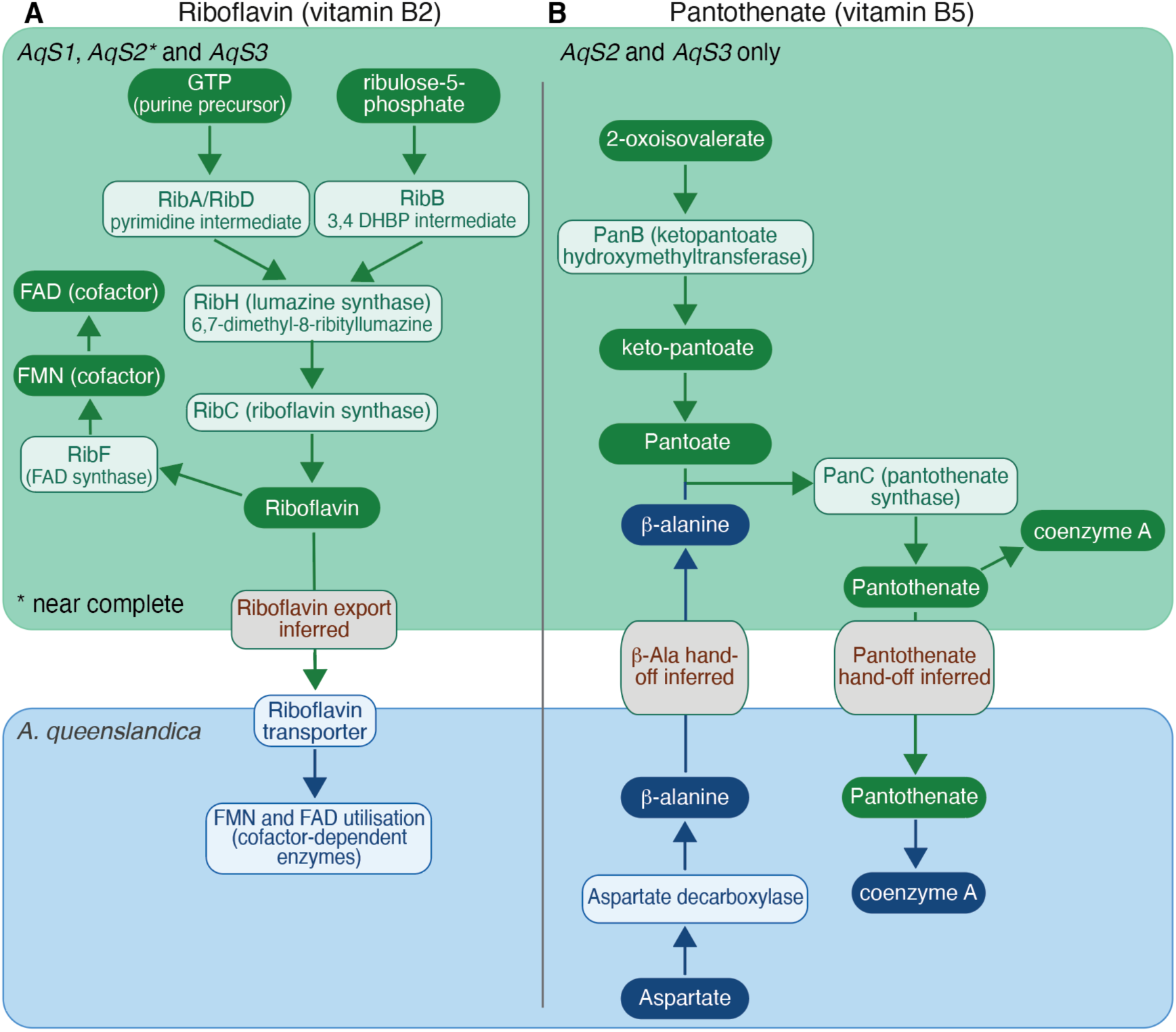
Riboflavin and pantothenate biosynthesis in the *A. queenslandica* holobiont. Schematics summarising the synthesis of riboflavin (vitamin B2) (A, left) and pantothenate (vitamin B5) (B, right). **(A)** Precursors and enzymes necessary for riboflavin synthesis are complete and highly expressed in the symbionts *AqS1* and *AqS3* and near-complete in *AqS2* (Fig. S15), but absent in the host sponge. *A. queenslandica* has a riboflavin transporter, which appears to be the sole host-side entry point. Symbionts and sponge can biosynthesise riboflavin derived co-factors FMN and FAD. **(B)** Substrates for the synthesis of pantothenate, and pantoate and β-alanine, are synthesised in *AqS2* and *AqS3*, and *A. queenslandica*, respectively (Fig. S16). Pantothenate-derived coenzyme A can be synthesised in symbionts and sponge host. The reciprocal transfer of β-alanine from the host and pantothenate from the symbiont is inferred as specific transporters were not identified. Solid arrows, inferred directional biosynthesis flow; blue (host) and green (symbiont) blocks, complete and highly expressed (Q2-Q4) pathways; coral blocks, partial inferred; metabolites, dark coloured pills.

Pantothenate (vitamin B5) biosynthesis illustrated a finer-grained form of integration, incorporating within-pathway partitioning across partners (Fig. 4 and Fig. S16). The host encoded highly expressed enzymes producing β-alanine, while *AqS2* and *AqS3* encoded *panC* and upstream steps consistent with pantoate supply. Neither partner alone completes the pathway, instead both contributions are required, and exchange of a biosynthetic intermediate is implied. This within-pathway division of labour is consistent with metabolite exchange being integral to the holobiont function (West and Cooper 2016; Tsoi et al. 2018).

### Complementary inorganic nutrient assimilation links environmental substrates to host metabolism

Dissolved inorganic nitrogen, sulfur, and phosphorus assimilation each showed complementary functional distributions across partners, with symbionts consistently encoding the reactions that transform environmental substrates into forms the host can assimilate (Fig. 5, Tables S10-S12 and Figs S17-19).

**Fig. 5.**
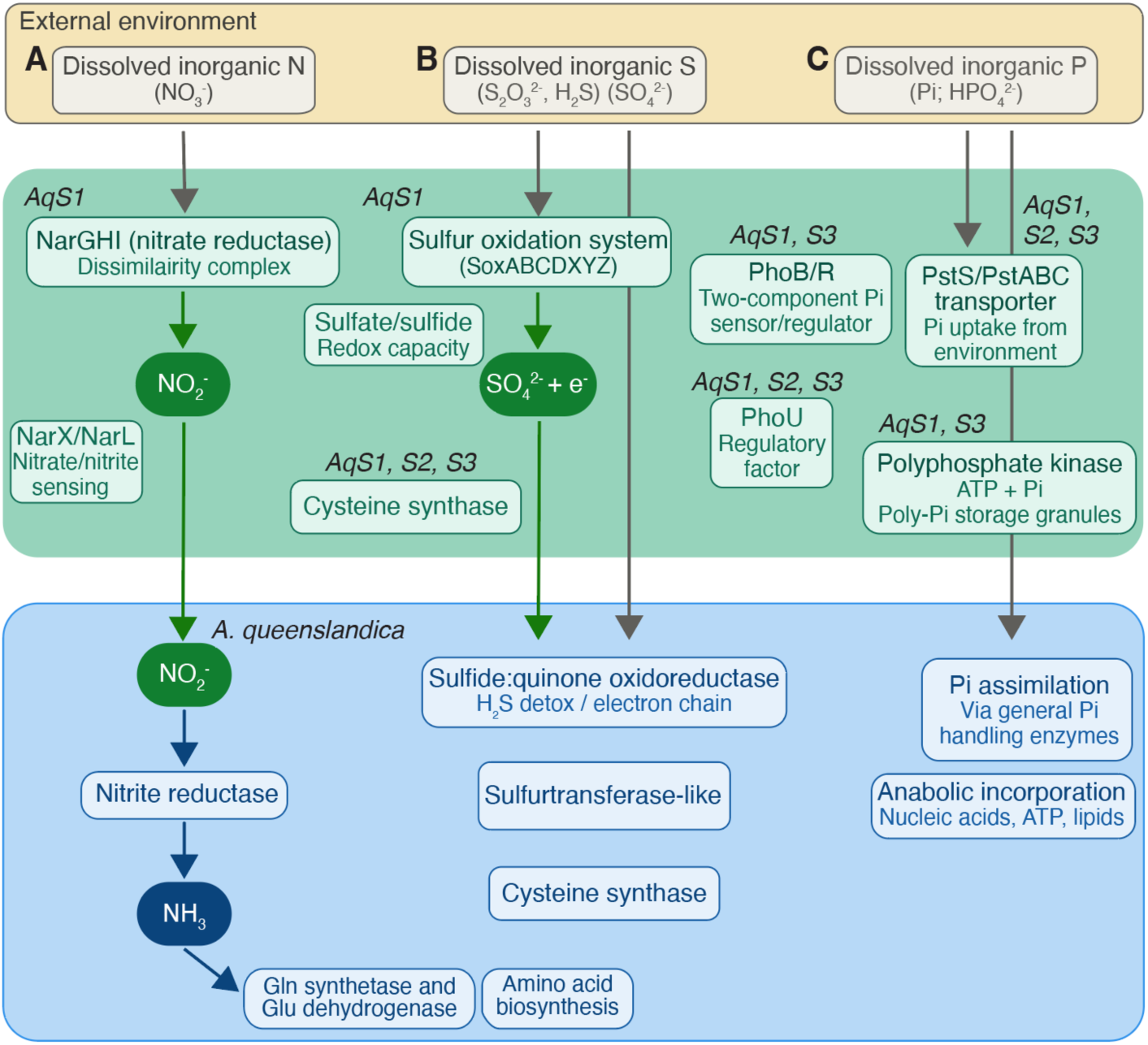
Schematics summarising the assimilation of dissolved inorganic nitrogen, sulfur, and phosphorus into the *A. queenslandica* holobiont. Inferred directional flow of nitrogen (A, left), sulfur (B, middle) and phosphorous (C, right) compounds between symbionts (*AqS1*, *AqS2* and *AqS3*) and *A. queenslandica* host, with source (environmental) inorganic compounds at the top. **(A)** *AqS1*’s nitrate reductase assimilation complex (NarGHI) reduces nitrate (NO_3_^-^) to nitrite (NO_2_^-^), and *A. queenslandica* reduces nitrite – some derived from exchange from *AqS1* - to ammonia via nitrite reductase. Glutamine (Gln) synthetase and glutamate (Glu) dehydrogenase incorporates ammonia into glutamine and glutamate, respectively, which contribute to downstream amino acid biosynthesis. Neither *AqS2* nor *AqS3* encode the nitrate reduction steps, and the sponge host lacks nitrate reductase. **(B)** Sulfur transformation is dominated by *AqS1*, which encodes and expresses the sulfur oxidation (SOX) system (SoxABCDXYZ components; Q2–Q4) for oxidation of environmental thiosulfate (S₂O₃²⁻) and sulfide (H₂S) to sulfate (SO₄²⁻), alongside broader sulfate–sulfide redox capacity. Reduced sulfur products are passed to the host for handling via SQOR, sulfurtransferase-like enzymes, and cysteine synthase. Cysteine synthesis is shared across all four partners. **(C)** The full Pho regulon cascade in *AqS1* and *AqS3* (PhoB/R → PstS/ABC → PhoU → PPK → polyP granules), with *AqS2* lacking all components except PstS and PhoU. See Figs 2 and 4 and Fig. S19 for details.

For nitrogen, *AqS1* encoded and expressed nitrate/nitrite sensing and nitrate reduction functions, while the host expressed nitrite reductase and downstream ammonia assimilation enzymes (glutamine synthetase and glutamate dehydrogenase). This is consistent with a sequential nitrate→nitrite→ammonia processing pathway in which *AqS1* performs the first reduction step and the host completes nitrogen incorporation into amino acids (Fig. 5 and Fig. S17). With *AqS1* performing the energetically demanding first reduction and the host completing nitrogen assimilation, this appears to be a functional partitioning across the host–symbiont interface. Sulfur metabolism was dominated by *AqS1*, which expressed genes consistent with thiosulfate and sulfide oxidation via SOX components, as well as broader sulfate–sulfide redox capacity (Fig. 5 and Fig. S18). Reduced sulfur compounds generated or transiting through *AqS1* are handled by the host via sulfide:quinone oxidoreductase (SQOR), which couples sulfide detoxification to the electron transport chain, and sulfurtransferase-like enzymes. Cysteine synthase genes were expressed across all four holobiont partners, consistent with cysteine biosynthesis being a shared metabolic function. *AqS2* and *AqS3* contribute to cysteine synthesis but do not encode the SOX system or dominate the sulfur oxidation capacity of the holobiont.

Phosphate acquisition and storage capacity were concentrated in *AqS1* and *AqS3*, with both encoding and expressing genes for Pho regulon components (PhoBR two-component phosphate sensor/regulator; high-affinity PstS phosphate-binding protein; PstABC ABC transporter for phosphate uptake from the environment; regulatory coupling factor PhoU; and polyphosphate kinase (PPK) for ATP-dependent synthesis and accumulation of intracellular polyphosphate granules), consistent with regulated phosphate uptake and polyphosphate accumulation (Fig. 5 and Fig. S19). The concentration of Pho regulon components in *AqS1* and *AqS3* positions these symbionts as the principal mediators of environmental phosphate acquisition for the holobiont under oligotrophic reef conditions.

*AqS2* lacks the Pho regulon, PstABC, and PPK, representing a notable functional distinction among the three symbionts and indicating specialisation of phosphorus acquisition within the consortium. *A. queenslandica* encodes general phosphate-handling enzymes and assimilates phosphate into biomass through nucleic acid, ATP, and lipid biosynthesis.

Together, these nutrient assimilation reconstructions identify multiple pathways in which symbiont-encoded transformations and expressed transporters connect dissolved environmental substrates to host assimilation capacity, extending the exchange-oriented architecture observed for carbon, amino acids, and vitamins into the domain of inorganic nutrient cycling.

### Membrane transport enrichment as the unifying axis of holobiont integration

Across carbon utilisation, amino acid biosynthesis, vitamin and cofactor availability, and inorganic nutrient assimilation, the three symbionts consistently contribute disproportionate metabolic breadth and encode the majority of membrane transport machinery in the holobiont (Figs 1 and 6 and Tables S6 and S7). In each case, symbiont biosynthetic or transformation capacity is paired with expressed membrane transporter repertoires on both symbiont and host sides, consistent with directed metabolite exchange rather than metabolic self-sufficiency. Symbiont membrane transporter genes are not distributed randomly across substrate classes but are concentrated in the uptake and export functions that complement the metabolism of the sugars, amino acids, inorganic ions, and vitamins identified in this study. The partner-specific functional contributions of the host and of the symbionts *AqS1*, *AqS2*, and *AqS3* are summarised in Figure 6.

**Fig. 6.**
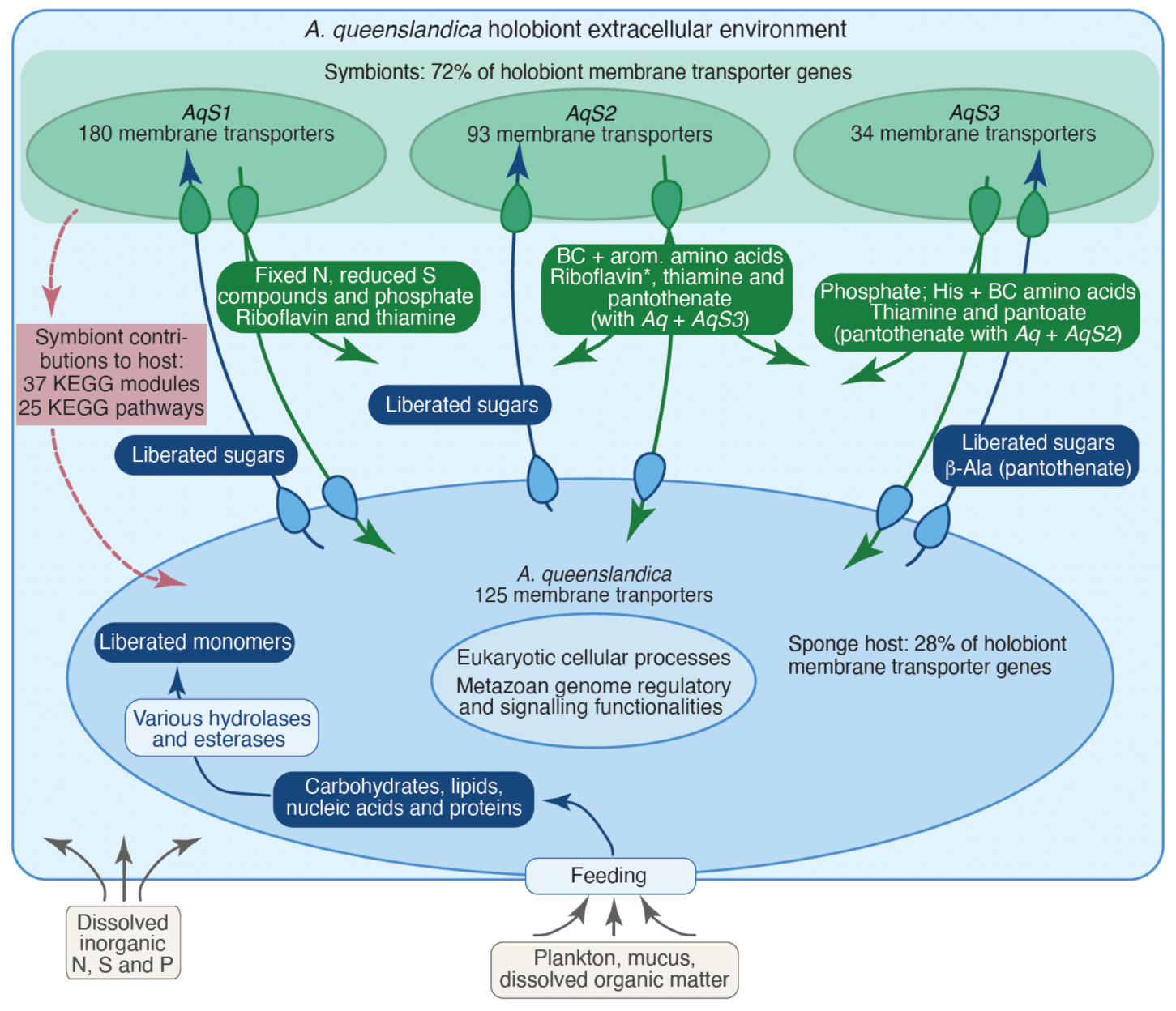
Transport-enabled metabolic expansion in the *Amphimedon queenslandica* holobiont. A synthesis schematic summarising the exchange-orientated functional architecture of the *A. queenslandica* holobiont, integrating the partner-resolved findings presented in Figures 2–5. All exchange flows are inferred from partner-resolved KEGG gene content and adult holotranscriptome expression (mean expression quartile ≥Q2 across nine holotranscriptomes) (Tables S10 and S11). The host sponge cell (blue) is shown at the bottom and the three vertically inherited gammaproteobacterial symbionts, *AqS1*, *AqS2* and *AqS3* (green) are at the top, with all partners enclosed in a common extracellular environment (light blue) containing predominantly host-derived extracellular biomolecules. The inferred directional movement of nutrients, monomers and other biomolecules into and out of cells are listed in dark coloured pills (symbionts, dark green; host, dark blue) with arrows that are coloured based on cell of origin (for vitamins: * riboflavin synthesis complete in *AqS2* but one step was expressed below the Q2 threshold). Membrane transporters on cell surfaces are represented by teardrops that are oriented to show general import and export functions in host and symbionts. Red box and dashed arrows show number of KEGG modules and pathways that are incomplete in *A. queenslandica* but are completed through symbiont contributions (Table 2). Food and inorganic nutrient inputs from the external environment are shown outside the holobiont at the bottom in grey. Numbers of membrane transporters encoded in host and symbiont genomes are shown. β-Ala, β-alanine; BC, branched-chain; His, histidine; N, nitrogen; P, phosphorous; S, sulfur.

## Discussion

### A small symbiont consortium disproportionately shapes holobiont metabolism through exchange-oriented specialisation

The central finding of this study is not that symbionts contribute to *A. queenslandica* metabolism – that much has been established already for sponges (Slaby et al. 2017; Gantt et al. 2019; Engelberts et al. 2020; Song et al. 2021; Hudspith et al. 2021; Moeller et al. 2023) – but that three vertically inherited bacteria disproportionately expand holobiont metabolic capacity through a functional architecture centred on membrane transport and metabolite exchange (Ren and Paulsen 2005; Clarke et al. 2014). Despite representing only ∼20% of KEGG-annotated hologenome genes, *AqS1*, *AqS2*, and *AqS3* encode nearly half of all metabolic genes and almost three-quarters of membrane transport genes. This imbalance reflects functional specialisation (West and Cooper 2016; Tsoi et al. 2018) in which symbionts contribute metabolic breadth and exchange capacity while the host provides the regulatory, signalling, and multicellular physiological context into which symbiont-derived metabolites are integrated. The result is a holobiont whose collective metabolic capacity substantially exceeds what host or symbionts could sustain independently, assembled from just three microbial partners.

Importantly, symbiont contributions are not confined to peripheral or conditionally expressed pathways. They span central carbon metabolism, amino acid biosynthesis, vitamin and cofactor production, and inorganic nutrient assimilation, with associated genes expressed at moderate to high levels in all adult holotranscriptomes. That genome-predicted capacities are transcriptionally active under both natural and aquaria-maintained conditions substantially strengthens the inference that symbionts are functional metabolic partners rather than metabolically capable but physiologically disengaged associates.

The mechanism connecting symbiont metabolic breadth to host physiology is membrane transport. Across every pathway examined, symbiont biosynthetic or transformation capacity is paired with enriched membrane transporter repertoires on both symbiont and host sides, consistent with directed metabolite flow. Symbionts encode the dominant share of sugar, amino acid, inorganic ion, and vitamin transporters in the holobiont, while the host expresses uptake systems matched to the specific compounds that symbionts produce. This reciprocal transport architecture distinguishes the present study, which infers metabolic integration from genome content combined with transcriptional activity, from those in which microbial metabolic capacity is inferred from biosynthetic gene content alone (Slaby et al. 2017; Engelberts et al. 2020; Gantt et al. 2019). Membrane transporter repertoire has often been under analysed in holobiont research. Our results argue that transporter repertoires deserve systematic attention alongside biosynthetic pathway reconstruction in any system where metabolic integration between host and microbe is proposed.

### Implications for sponge ecology and biogeochemical cycling

The metabolic architecture revealed here connects directly to the ecological role of sponges as major benthic–pelagic couplers in oligotrophic reef systems (de Goeij et al. 2013; Maldonado 2015; Pawlik et al. 2016; Pita et al. 2018). Shared heterotrophic carbon metabolism across all four partners supports efficient assimilation of dissolved organic matter, while symbiont contributions to amino acid, vitamin, and cofactor biosynthesis extend the nutritional repertoire of the holobiont beyond what the host genome alone could support. Complementary pathways for nitrogen, sulfur, and phosphorus assimilation position symbionts as important mediators of inorganic nutrient transformation and retention within sponge biomass (Zhang et al. 2019).

The broad transcriptional activity of symbiont-enriched metabolic and membrane transporter genes in adult holobionts is consistent with these pathways and functionalities being operational under normal physiological conditions, supporting a model in which vertically inherited symbionts enhance sponge performance in nutrient-limited environments by expanding metabolic options, facilitating exchange, and buffering nutrient availability (Pita et al. 2018; Glasl et al. 2024). At the reef scale, this would amplify the well-documented role of sponges in benthic nutrient retention and recycling (de Goeij et al. 2013; Maldonado 2015; Pawlik et al. 2016; Pita et al. 2018), not merely through filter feeding, but through symbiont-mediated transformation of dissolved inorganic nutrients into holobiont biomass (Fiore et al. 2013; Gantt et al. 2019; Zhang et al. 2019; Glasl et al. 2024).

### Low microbiome complexity enables functional attribution

A persistent challenge in holobiont and microbiome research is disentangling the contributions of individual microbial partners in complex, strain-rich communities. This is particularly challenging in non-model, yet ecologically important, holobionts (Pita et al. 2018; Engelberts et al. 2020; Carrier et al. 2022). Metagenomes from diverse holobiotic communities can carry assembly errors, strain-level ambiguity, and incomplete coverage, making confident functional attribution difficult. The *A. queenslandica* holobiont largely circumvents these problems. Its microbiome is dominated by three proteobacterial symbionts that are vertically transmitted, seasonally stable (Fieth et al. 2016; Gauthier et al. 2016), and represented here by improved partner-specific genome assemblies with matched holotranscriptomes, fitting a broader pattern in which vertical transmission can stabilise sponge symbiont composition and function across reproductive stages and generations (Schmitt et al. 2008; Carrier et al. 2022; Carrier et al. 2023; Glasl et al. 2024). This simplicity allows metabolic capacities to be assigned unambiguously to individual partners and compared directly with the host.

This partner-level resolution allows for tractable mechanistic inferences. For instance, the disproportionate enrichment of metabolic and membrane transporter genes in symbionts, the complementary distribution of biosynthetic and assimilation functions across partners, and the within-pathway partitioning observed for pantothenate biosynthesis would be difficult or impossible to resolve in a more complex microbiome. We acknowledge that transcript abundance may not equate directly to enzymatic activity or metabolite flux, and that some symbiont pathways may be condition-specific or regulated post-transcriptionally in ways not captured here (Nie et al. 2006; Dressaire et al. 2009; Mehmeti et al. 2012). Nonetheless, the *A. queenslandica* holobiont illustrates the value of low-complexity holobionts as tractable models for understanding principles of metabolic integration, and the specific role of membrane transport within that integration, that may operate more broadly across animal–microbe symbioses.

### A model for transport-enabled metabolic expansion in animal–microbe symbiosis

The *A. queenslandica* holobiont reveals how a small number of vertically inherited microbial symbionts can disproportionately expand host metabolic capacity not through numerical or genomic dominance, but through specialisation in metabolic breadth and membrane transport. This exchange-oriented capacity allows symbionts to act as metabolic hubs that function in transforming substrates, synthesising compounds the host cannot make, and redistributing metabolites through enriched transporter repertoires (Ren and Paulsen 2005; Clarke et al. 2014). The sponge host integrates these inputs into multicellular physiology and biomass, consistent with stable-isotope and NanoSIMS evidence that metabolites and nutrients can be exchanged between sponge host cells and microbial partners (Rix et al. 2020; Hudspith et al. 2021; Moeller et al. 2023). Symbionts here do not simply duplicate host pathways, but instead provide qualitatively different functional capacities that the host genome has not retained or never possessed. Membrane transporters make this expansion and partitioning functional by directly connecting symbiont metabolic breadth to host physiology (Clarke et al. 2014; Moeller et al. 2023) across a physical interface that separates extracellular bacteria from sponge cells.

More broadly, these findings suggest that transporter repertoire enrichment, alongside biosynthetic pathway complementation, should be a primary target of analysis in any proposed nutritional or metabolic symbiosis. By resolving these relationships at partner-specific resolution in a low-complexity system, the *A. queenslandica* holobiont provides a valuable system for understanding how microbial symbionts shape animal metabolism, ecology, and evolution.

**Supplementary Information** 12 Supplementary tables and 19 Supplementary figures accompany this paper.

## Acknowledgements

The authors thank staff at the Heron Island Research Station for logistical assistance during field collections, and Nick Rhodes at QCIF for management of high-performance computing facilities.

## Author contributions

Xueyan Xiang: conceptualisation, methodology, visualisation, resources, Writing – original draft preparation; Eve Maunders: methodology, visualisation, Writing – original draft preparation; Bernard M Degnan: conceptualisation, methodology, funding acquisition, resources, project administration, supervision, Writing – original draft preparation; Sandie M Degnan: conceptualisation, methodology, funding acquisition, resources, project administration, supervision, Writing – original draft preparation.

## Data availability

New *A. queenslandica*, *AqS1*, *AqS2* and *AqS3* genome assemblies and annotations are available in NCBI (BioProject PRJNA6686600).

## Ethics approval

Adult *Amphimedon queenslandica* were collected from Heron Island Reef, under GBRMPA permit G16/38120.1. No other permits were required for this research.

## References

Anders S, Pyl PT, Huber W (2015) HTSeq--a Python framework to work with high-throughput sequencing data. Bioinformatics 31:166–169. 10.1093/bioinformatics/btu638

Aramaki T, Blanc-Mathieu R, Endo H, Ohkubo K, Kanehisa M, Goto S, Ogata H (2020) KofamKOALA: KEGG ortholog assignment based on profile HMM and adaptive score threshold. Bioinformatics 36:2251–2252. 10.1093/bioinformatics/btz859

Bayer K, Jahn MT, Slaby BM, Moitinho-Silva L, Hentschel U (2018) Marine sponges as *Chloroflexi* hot spots: genomic insights and high-resolution visualization of an abundant and diverse symbiotic clade. mSystems 3:e00150–18. 10.1128/mSystems.00150-18

Bell JJ (2008) The functional roles of marine sponges. Estuar Coast Shelf Sci 79:341–353. 10.1016/j.ecss.2008.05.002

Bolger AM, Lohse M, Usadel B (2014) Trimmomatic: a flexible trimmer for Illumina sequence data. Bioinformatics 30:2114–2120. 10.1093/bioinformatics/btu170

Burgin J, Allen B, Wilkinson DJ, Beracochea M, Escobar-Zepeda A, Korobeynikov A, Richardson L, Colwell LJ (2023) MGnify: the microbiome sequence data analysis resource in 2023. Nucleic Acids Res 51:D753–D759. 10.1093/nar/gkac1080

Carrier TJ, Maldonado M, Schmittmann L, Pita L, Bosch TCG, Hentschel U (2022) Symbiont transmission in marine sponges: reproduction, development, and metamorphosis. BMC Biol 20:132. 10.1186/s12915-022-01291-6

Carrier TJ, Schmittmann L, Jung S, Pita L, Hentschel U (2023) Maternal provisioning of an obligate symbiont in a sponge. Ecol Evol 13:e10012. 10.1002/ece3.10012

Clarke VC, Loughlin PC, Day DA, Smith PMC (2014) Transport processes of the legume symbiosome membrane. Front Plant Sci 5:699. 10.3389/fpls.2014.00699

Conesa A, Götz S (2008) Blast2GO: A comprehensive suite for functional analysis in plant genomics. Int J Plant Genomics 2008:619832. 10.1155/2008/619832

de Goeij JM, van Oevelen D, Vermeij MJA, Osinga R, Middelburg JJ, de Goeij AFPM, Admiraal W (2013) Surviving in a marine desert: the sponge loop retains resources within coral reefs. Science 342:108–110. 10.1126/science.1241981

Dressaire C, Gitton C, Loubière P, Monnet V, Queinnec I, Cocaign-Bousquet M (2009) Transcriptome and proteome exploration to model translation efficiency and protein stability in *Lactococcus lactis*. PLoS Comput Biol 5:e1000606. 10.1371/journal.pcbi.1000606

Engelberts JP, Robbins SJ, de Goeij JM, Aranda M, Bell SC, Webster NS (2020) Characterization of a sponge microbiome using an integrative genome-centric approach. ISME J 14:1100–1110. 10.1038/s41396-020-0591-9

Fernandez-Valverde SL, Calcino AD, Degnan BM (2015) Deep developmental transcriptome sequencing uncovers numerous new genes and enhances gene annotation in the sponge *Amphimedon queenslandica*. BMC Genomics 16:387. 10.1186/s12864-015-1588-z

Fieth RA, Gauthier MEA, Bayes J, Green KM, Degnan SM (2016) Ontogenetic changes in the bacterial symbiont community of the tropical demosponge *Amphimedon queenslandica*: metamorphosis is a new beginning. Front Mar Sci 3:228. 10.3389/fmars.2016.00228

Fiore CL, Baker DM, Lesser MP (2013) Nitrogen biogeochemistry in the Caribbean sponge, *Xestospongia muta*: a source or sink of dissolved inorganic nitrogen? PLoS One 8:e72961. 10.1371/journal.pone.0072961

Fiore CL, Labrie M, Jarettt JK, Lesser MP (2015) Transcriptional activity of the giant barrel sponge, *Xestospongia muta* holobiont: molecular evidence for metabolic interchange. Front Microbiol 6:364. 10.3389/fmicb.2015.00364

Folkers M, Rombouts T (2020) Sponges revealed: a synthesis of their overlooked ecological functions within aquatic ecosystems. YOUMARES 9 - The Oceans: Our Research, Our Future:181–193. 10.1007/978-3-030-20389-4_9

Gantt SE, McMurray SE, Stubler AD, Finelli CM, Pawlik JR, Erwin PM (2019) Testing the relationship between microbiome composition and flux of carbon and nutrients in Caribbean coral reef sponges. Microbiome 7:124. 10.1186/s40168-019-0739-x

Gauthier MEA, Watson JR, Degnan SM (2016) Draft genomes shed light on the dual bacterial symbiosis that dominates the microbiome of the coral reef sponge *Amphimedon queenslandica*. Front Mar Sci 3:196. 10.3389/fmars.2016.00196

Glasl B, Luter HM, Damjanovic K, Kitzinger K, Mueller AJ, Mahler L, Engelberts JP, Rix L, Osvatic JT, Hausmann B, Séneca J, Daims H, Pjevac P, Wagner M (2024) Co-occurring nitrifying symbiont lineages are vertically inherited and widespread in marine sponges. ISME J 18:wrae069. 10.1093/ismejo/wrae069

Hudspith M, Rix L, Achlatis M, Bougoure J, Guagliardo P, Clode PL, Webster NS, Muyzer G, Pernice M, de Goeij JM (2021) Subcellular view of host–microbiome nutrient exchange in sponges: insights into the ecological success of an early metazoan–microbe symbiosis. Microbiome 9:37. 10.1186/s40168-020-00984-w

Kamke J, Sczyrba A, Ivanova N, Schwientek P, Rinke C, Mavromatis K, Woyke T, Hentschel U (2013) Single-cell genomics reveals complex carbohydrate degradation patterns in poribacterial symbionts of marine sponges. ISME J 7:2287–2300. 10.1038/ismej.2013.111

Kanehisa M, Sato Y, Morishima K (2016) BlastKOALA and GhostKOALA: KEGG tools for functional characterization of genome and metagenome sequences. J Mol Biol 428:726–731. 10.1016/j.jmb.2015.11.006

Kim D, Langmead B, Salzberg SL (2015) HISAT: a fast spliced aligner with low memory requirements. Nat Methods 12:357–360. 10.1038/nmeth.3317

Leys SP, Yahel G, Reidenbach MA, Tunnicliffe V, Shavit U, Reiswig HM (2011) The sponge pump: the role of current induced flow in the design of the sponge body plan. PLoS One 6:e27787. 10.1371/journal.pone.0027787

Liao H, Myung S, Zhang Y-HP (2012) One-step purification and immobilization of thermophilic polyphosphate glucokinase from *Thermobifida fusca* YX: glucose-6-phosphate generation without ATP. Appl Microbiol Biotechnol 93:1109–1117. 10.1007/s00253-011-3458-1

Li H, Durbin R (2009) Fast and accurate short read alignment with Burrows–Wheeler transform. Bioinformatics 25:1754–1760. 10.1093/bioinformatics/btp324

Li Z, Sun W, Zhang F, He L, Loganathan K (2015) Actinomycetes from the South China Sea sponges: isolation, diversity and potential for aromatic polyketides discovery. Front Microbiol 6:1048. 10.3389/fmicb.2015.01048

Maldonado M (2015) Sponge waste that fuels marine oligotrophic food webs: a re-assessment of its origin and nature. Mar Ecol 37:477–491. 10.1111/maec.12256

Mehmeti, et al. 2012 Mehmeti I, Faergestad EM, Bekker M, Snipen L, Nes IF, Holo H (2012) Growth rate-dependent control in Enterococcus faecalis: effects on the transcriptome and proteome, and strong regulation of lactate dehydrogenase. Appl Environ Microbiol 78:170–176. 10.1128/AEM.06604-11

Moeller FU, Herbold CW, Schintlmeister A, Mooshammer M, Motti C, Glasl B, Kitzinger K, Behnam F, Watzka M, Schweder T, Albertsen M, Richter A, Webster NS, Wagner M (2023) Taurine as a key intermediate for host-symbiont interaction in the tropical sponge *Lanthella basta*. ISME J 17:1208–1223. 10.1038/s41396-023-01420-1

Moitinho-Silva L, Nielsen S, Amir A et al. (2017) The sponge microbiome project. GigaScience 6:gix077. 10.1093/gigascience/gix077

Nie, et al. 2006 Nie L, Wu G, Zhang W (2006) Correlation between mRNA and protein abundance in Desulfovibrio vulgaris: a multiple regression to identify sources of variations. Biochem Biophys Res Commun 339:603–610. 10.1016/j.bbrc.2005.11.055

O’Brien PA, Tan S, Yang C, Frade PR, Andreakis N, Smith HA, Miller DJ, Webster NS, Zhang G, Bourne DG (2020) Diverse coral reef invertebrates exhibit patterns of phylosymbiosis. ISME J 14:2211–2222. 10.1038/s41396-020-0671-x

Pawlik JR, Burkepile DE, Thurber RV (2016) A vicious circle? Altered carbon and nutrient cycling may explain the low resilience of Caribbean coral reefs. BioScience 66:470–476. 10.1093/biosci/biw047

Pita L, Rix L, Slaby BM, Franke A, Hentschel U (2018) The sponge holobiont in a changing ocean: from microbes to ecosystems. Microbiome 6:46. 10.1186/s40168-018-0428-1

Pracana R, Priyam A, Levantis I, Nichols RA, Wurm Y (2017) The fire ant social chromosome supergene variant Sb shows low diversity but high divergence from SB. Mol Ecol 26:2864–2879. 10.1111/mec.14054

Putnam NH, O’Connell BL, Stites JC et al (2016) Chromosome-scale shotgun assembly using an in vitro method for long-range linkage. Genome Res 26:342–350. 10.1101/gr.193474.115

Quinlan AR, Hall IM (2010) BEDTools: a flexible suite of utilities for comparing genomic features. Bioinformatics 26:841–842. 10.1093/bioinformatics/btq033

Ren Q, Paulsen IT (2005) Comparative analyses of fundamental differences in membrane transport capabilities in prokaryotes and eukaryotes. PLoS Comput Biol 1:e27. 10.1371/journal.pcbi.0010027

Rix L, de Goeij JM, Mueller CE, Struck U, Middelburg JJ, van Duyl FC, Al-Horani FA, Wild C, Naumann MS, van Oevelen D (2016) Coral mucus fuels the sponge loop in warm- and cold-water coral reef ecosystems. Sci Rep 6:18715. 10.1038/srep18715

Rix L, de Goeij JM, van Oevelen D, Struck U, Al-Horani FA, Wild C, Naumann MS (2017) Differential recycling of coral and algal dissolved organic matter via the sponge loop. Funct Ecol 31:778–789. 10.1111/1365-2435.12758

Rix L, Ribes M, Coma R, Jahn MT, de Goeij JM, van Oevelen D, Escrig S, Meibom A, Hentschel U (2020) Heterotrophy in the earliest gut: a single-cell view of heterotrophic carbon and nitrogen assimilation in sponge-microbe symbioses. ISME J 14:2554–2567. 10.1038/s41396-020-0706-3

Schmitt S, Angermeier H, Schiller R, Lindquist N, Hentschel U (2008) Molecular microbial diversity survey of sponge reproductive stages and mechanistic insights into vertical transmission of microbial symbionts. Appl Environ Microbiol 74:7694–7708. 10.1128/aem.00878-08

Seemann T (2014) Prokka: rapid prokaryotic genome annotation. Bioinformatics 30:2068–2069. 10.1093/bioinformatics/btu153

Simão FA, Waterhouse RM, Ioannidis P, Kriventseva EV, Zdobnov EM (2015) BUSCO: assessing genome assembly and annotation completeness with single-copy orthologs. Bioinformatics 31:3210–3212. 10.1093/bioinformatics/btv351

Slaby BM, Hackl T, Horn H, Bayer K, Hentschel U (2017) Metagenomic binning of a marine sponge microbiome reveals unity in defense but metabolic specialization. ISME J 11:2465–2478. 10.1038/ismej.2017.101

Song H, Hewitt OH, Degnan SM (2021) Arginine biosynthesis by a bacterial symbiont enables nitric oxide production and facilitates larval settlement in the marine-sponge host. Curr Biol 31:433–437. 10.1016/j.cub.2020.10.051

Srivastava M, Simakov O, Chapman J et al (2010) The *Amphimedon queenslandica* genome and the evolution of animal complexity. Nature 466:720–726. 10.1038/nature09201

Steinert G, Busch K, Bayer K et al (2020) Compositional and quantitative insights into bacterial and archaeal communities of South Pacific deep-sea sponges (Demospongiae and Hexactinellida). Front Microbiol 11:716. 10.3389/fmicb.2020.00716

Taylor JA, Palladino G, Wemheuer B, Steinert G, Sipkema D, Williams TJ, Thomas T (2021) Phylogeny resolved, metabolism revealed: functional radiation within a widespread and divergent clade of sponge symbionts. ISME J 15:503–519. 10.1038/s41396-020-00791-z

Thomas T, Rusch D, DeMaere MZ et al (2010) Functional genomic signatures of sponge bacteria reveal unique and shared features of symbiosis. ISME J 4:1557–1567. 10.1038/ismej.2010.74

Tsoi R, Wu F, Zhang C, Bewick S, Karig D, You L (2018) Metabolic division of labor in microbial systems. Proc Natl Acad Sci USA 115:2526–2531. 10.1073/pnas.1716888115

Wagner GP, Kin K, Lynch VJ (2012) Measurement of mRNA abundance using RNA-seq data: RPKM measure is inconsistent among samples. Theory Biosci 131:281–285. 10.1007/s12064-012-0162-3

Webster NS, Thomas T (2016) The sponge hologenome. mBio 7:e00135–16. 10.1128/mBio.00135-16

West SA, Cooper GA (2016) Division of labour in microorganisms: an evolutionary perspective. Nat Rev Microbiol 14:716–723. 10.1038/nrmicro.2016.111

Xiang X, Poli D, Degnan BM, Degnan SM (2022) Ribosomal RNA-depletion provides an efficient method for successful dual RNA-Seq expression profiling of a marine sponge holobiont. Mar Biotechnol 24:722–732. 10.1007/s10126-022-10138-8

Yang B, Yuen-Simović B, Yuan H, Degnan BM, Degnan SM (2026) Early transcription factor activation distinguishes symbiotic from non-symbiotic bacteria during microbiome processing in a sponge. ISME J 20:wrag150. 10.1093/ismejo/wrag150

Yu Z, Meng L, Nguyen CH, Mamitsuka H, Kanehisa M, Ogata H (2026) DeepKOALA: a fast and accurate deep learning framework for KEGG Orthology assignment. bioRxiv. 10.64898/2026.01.07.698072

Zhang F, Jonas L, Lin H, Hill RT (2019) Microbially mediated nutrient cycles in marine sponges. FEMS Microbiol Ecol 95:fiz155. 10.1093/femsec/fiz155

