## Supplemental figures S1-S19 for "A partner-resolved marine sponge hologenome reveals that three bacterial symbionts disproportionately expand holobiont metabolism via enriched membrane transporter repertoires"

**Supplementary figure legends**

**Fig. S1. Glycolysis/ gluconeogenesis pathway.** KEGG Orthology (KO) assigned host (*A. queenslandica*; *Aq*) and symbiont (*AqS1*, *AqS2* and *AqS3*) genes mapped onto KEGG modules and pathways. The EC code is designated in each enzyme block. The four sub-blocks in each enzyme block from left to right represent the expression levels in *A. queenslandica*, *AqS1*, *AqS2* and *AqS3*. The colour reflects the gene's relative expression within each organism (i.e. not the holobiont), with dark blue (4) and light yellow (0) corresponding to the most highly and lowly expressed genes, respectively. Genes expressed above 2 are in the top two quartiles.

**Fig. S2. Summary of glycolysis and glycolytic enzymes in the holobiont.** The EC codes are sub-blocked as per Fig. S1, with only expressed genes colour-coded.

**Fig. S3. Citrate (TCA) cycle.** See Fig. S1 legend for explanation of how the pathways is annotated.

**Fig. S4. Pentose phosphate pathway.** See Fig. S1 legend for explanation of how the pathway is annotated.

**Fig. S5. Summary of pentose phosphate pathway enzymes in the holobiont.** The EC codes are sub-blocked as per Fig. S1, with only expressed genes colour-coded.

**Fig. S6. Alanine, aspartate and glutamate metabolic pathways.** See Fig. S1 legend for explanation of how these pathways are annotated.

**Fig. S7. Arginine biosynthesis pathway.** See Fig. S1 legend for explanation of how the pathway is annotated.

**Fig. S8. Glycine, serine and threonine metabolic pathways.** See Fig. S1 legend for explanation of how these pathways are annotated.

**Fig. S9. Histidine metabolic pathway.** See Fig. S1 legend for explanation of how the pathway is annotated.

**Fig. S10. Lysine biosynthesis pathway.** See Fig. S1 legend for explanation of how the pathway is annotated.

**Fig. S11. Phenylalanine, tyrosine and tryptophan biosynthesis pathways.** See Fig. S1 legend for explanation of how the pathways are annotated.

**Fig. S12. Valine, leucine and isoleucine biosynthesis pathways.** See Fig. S1 legend for explanation of how the pathways are annotated.

**Fig. S13. Cysteine and methionine metabolic pathways.** See Fig. S1 legend for explanation of how these pathways are annotated.

**Fig. S14. Thiamine metabolic pathway.** See Fig. S1 legend for explanation of how the pathway is annotated.

**Fig. S15. Riboflavin metabolic pathway.** See Fig. S1 legend for explanation of how the pathway is annotated.

**Fig. S16. Pantothenate and CoA biosynthesis pathways.** See Fig. S1 legend for explanation of how the pathways are annotated.

**Fig. S17. Nitrogen metabolic pathway.** See Fig. S1 legend for explanation of how the pathway is annotated.

**Fig. S18. Sulfur metabolic pathway.** See Fig. S1 legend for explanation of how the pathway is annotated.

**Fig. S19. Polyphosphate accumulation in the *A. queenslandica* holobiont.** This schematic is a working model for polyphosphate storage in the holobiont and follows the labelling schemes as per Figs S2 and S4. PhoBR, the phosphate two-component regulator and sensor proteins; PstS, phosphate binding protein; PstABC, the phosphate ABC transporter; PhoU, the phosphate transport system regulatory protein; PPK, polyphosphate kinase; PolyP, polyphosphate. Figure adapted from Peng et al. (2017).

**Reference**

Peng YC, Lu C, Li G, Eichenbaum Z, Lu CD (2017) Induction of the pho regulon and polyphosphate synthesis against spermine stress in *Pseudomonas aeruginosa*. *Mol Micro* 104:1037–1051. https://doi.org/10.1111/mmi.13678

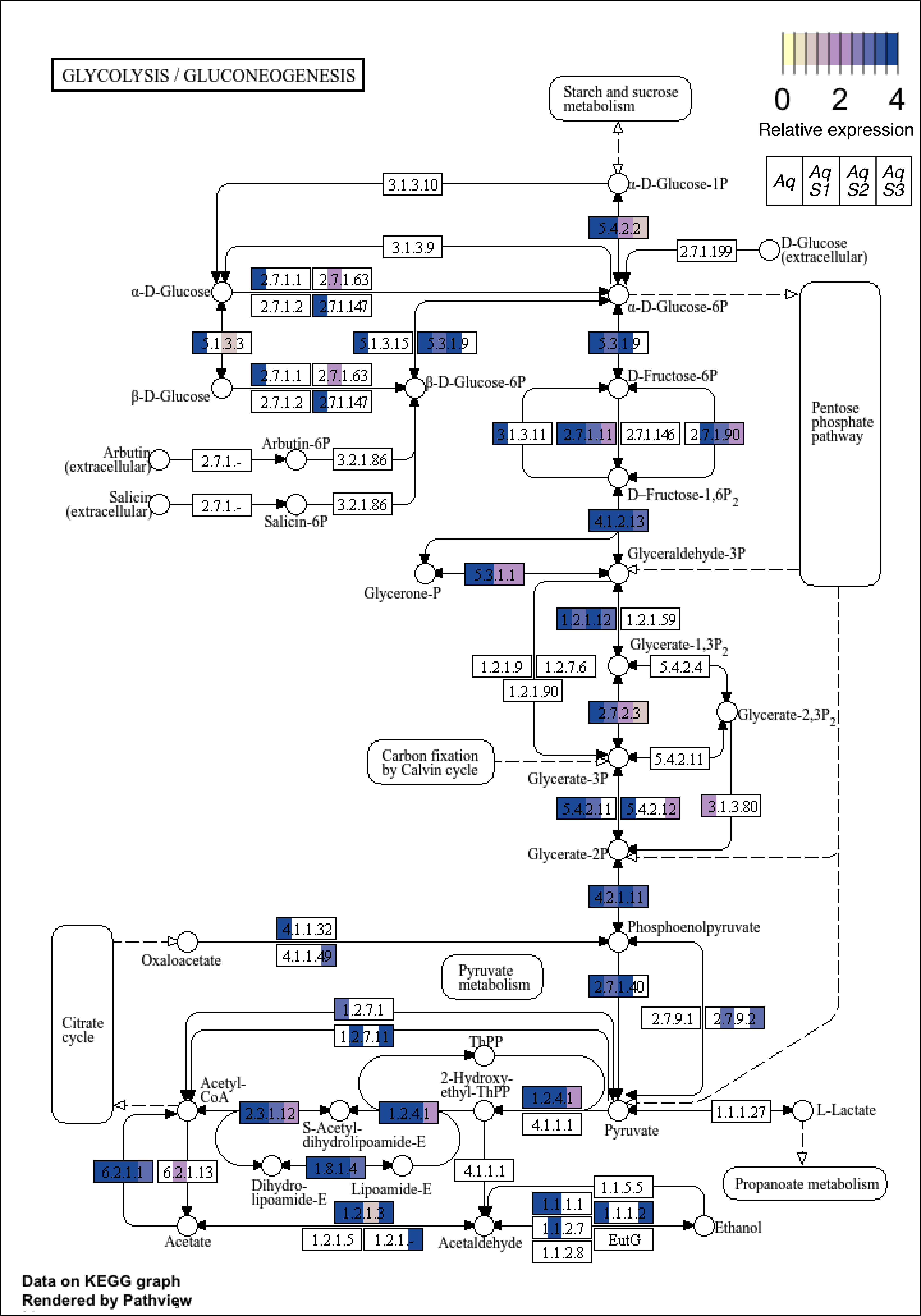

GLYCOLYSIS / GLUCONEOGENESIS
0 2 4
Relative expression
Aq
Aq S1
Aq S2
Aq S3
Starch and sucrose metabolism
α-D-Glucose-1P
3.1.3.10
5.4.2.2
3.1.3.9
2.7.1.199
D-Glucose (extracellular)
α-D-Glucose
2.7.1.1
2.7.1.63
2.7.1.2
2.7.1.147
α-D-Glucose-6P
Pentose phosphate pathway
5.1.3.3
5.1.3.15
5.3.1.9
5.3.1.9
β-D-Glucose
2.7.1.1
2.7.1.63
2.7.1.2
2.7.1.147
β-D-Glucose-6P
D-Fructose-6P
3.1.3.11
2.7.1.11
2.7.1.146
2.7.1.90
Arbutin (extracellular)
2.7.1.-
Arbutin-6P
3.2.1.86
Salicin (extracellular)
2.7.1.-
Salicin-6P
3.2.1.86
D-Fructose-1,6P2
4.1.2.13
Glyceraldehyde-3P
Glycerone-P
5.3.1.1
1.2.1.12
1.2.1.59
Glycerate-1,3P2
1.2.1.9
1.2.7.6
5.4.2.4
1.2.1.90
2.7.2.3
Glycerate-2,3P2
Carbon fixation by Calvin cycle
Glycerate-3P
5.4.2.11
5.4.2.11
5.4.2.12
3.1.3.80
Glycerate-2P
4.2.1.11
Phosphoenolpyruvate
4.1.1.32
4.1.1.49
Oxaloacetate
Citrate cycle
Pyruvate metabolism
2.7.1.40
2.7.9.1
2.7.9.2
1.2.7.1
1.2.7.11
ThPP
Acetyl-CoA
2-Hydroxy-ethyl-ThPP
1.2.4.1
2.3.1.12
S-Acetyl-dihydrolipoamide-E
1.2.4.1
4.1.1.1
Pyruvate
1.1.1.27
L-Lactate
6.2.1.1
6.2.1.13
Dihydro-lipoamide-E
1.8.1.4
Lipoamide-E
4.1.1.1
1.1.5.5
Propanoate metabolism
1.2.1.3
1.1.1.1
1.1.1.2
Acetate
1.2.1.5
1.2.1.-
Acetaldehyde
1.1.2.7
EutG
Ethanol
1.1.2.8
Data on KEGG graph
Rendered by Pathview

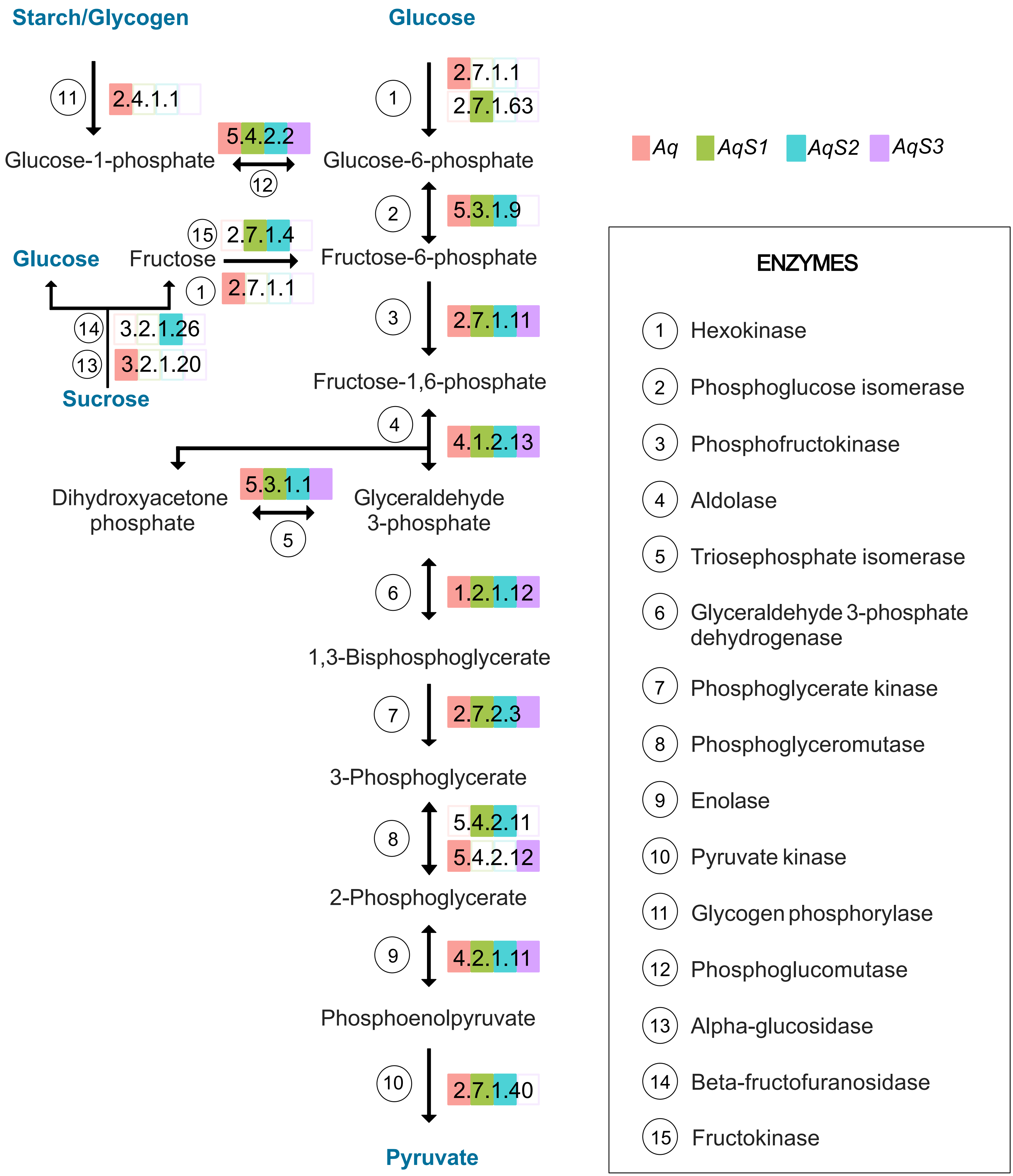
Starch/Glycogen
Glucose
11
2.4.1.1
1
2.7.1.1
2.7.1.63
Aq
AqS1
AqS2
AqS3
5.4.2.2
Glucose-1-phosphate
Glucose-6-phosphate
12
2
5.3.1.9
15
2.7.1.4
Glucose
Fructose
Fructose-6-phosphate
1
2.7.1.1
3
2.7.1.11
14
3.2.1.26
13
3.2.1.20
Sucrose
Fructose-1,6-phosphate
4
4.1.2.13
5.3.1.1
Dihydroxyacetone phosphate
Glyceraldehyde 3-phosphate
5
6
1.2.1.12
1,3-Bisphosphoglycerate
7
2.7.2.3
3-Phosphoglycerate
8
5.4.2.11
5.4.2.12
2-Phosphoglycerate
9
4.2.1.11
Phosphoenolpyruvate
10
2.7.1.40
Pyruvate
ENZYMES
1 Hexokinase
2 Phosphoglucose isomerase
3 Phosphofructokinase
4 Aldolase
5 Triosephosphate isomerase
6 Glyceraldehyde 3-phosphate dehydrogenase
7 Phosphoglycerate kinase
8 Phosphoglyceromutase
9 Enolase
10 Pyruvate kinase
11 Glycogen phosphorylase
12 Phosphoglucomutase
13 Alpha-glucosidase
14 Beta-fructofuranosidase
15 Fructokinase

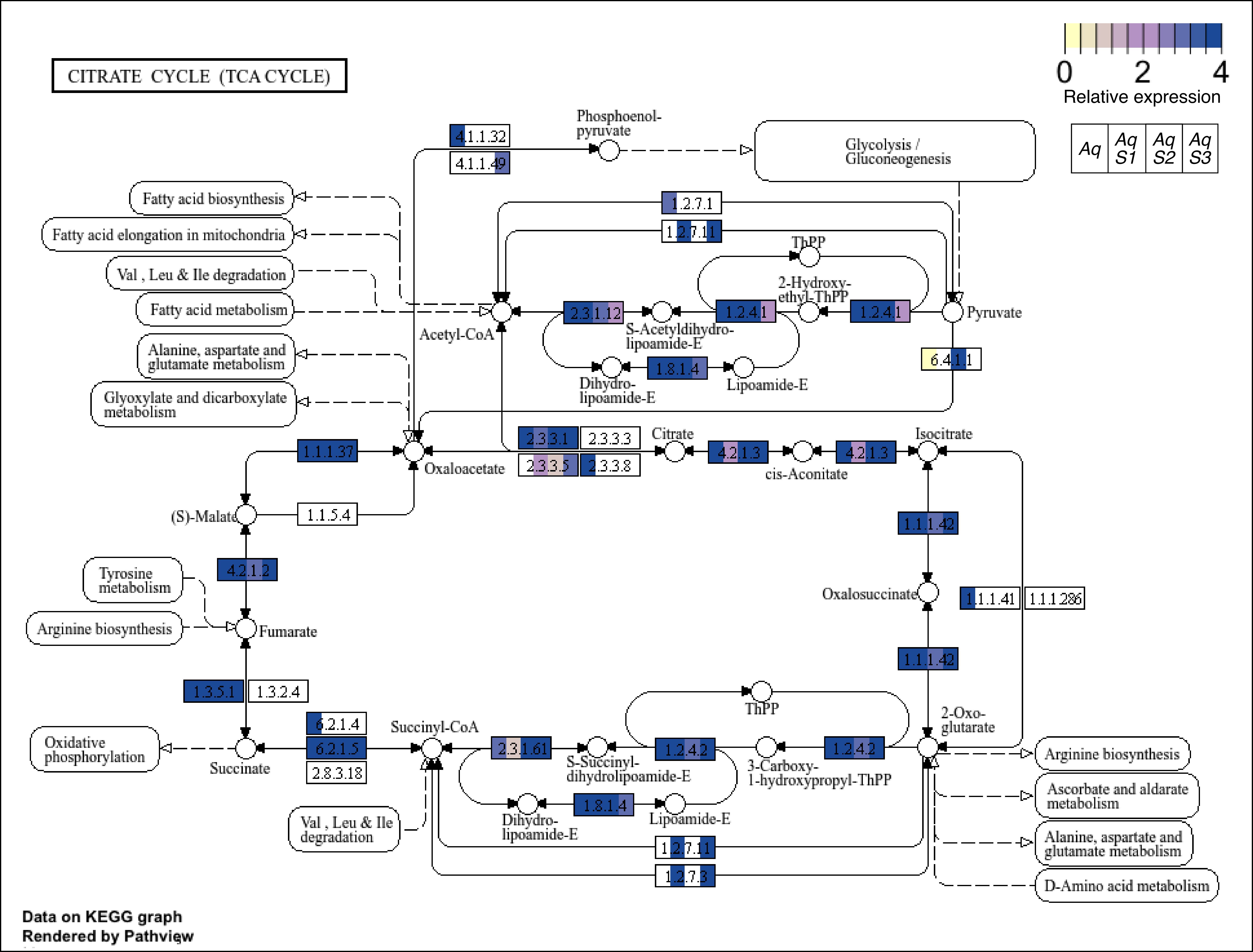

CITRATE CYCLE (TCA CYCLE)
0
2
4
Relative expression
Aq
Aq S1
Aq S2
Aq S3
Phosphoenol-pyruvate
Glycolysis / Gluconeogenesis
4.1.1.32
4.1.1.49
Fatty acid biosynthesis
Fatty acid elongation in mitochondria
Val , Leu & Ile degradation
Fatty acid metabolism
Alanine, aspartate and glutamate metabolism
Glyoxylate and dicarboxylate metabolism
1.2.7.1
1.2.7.11
ThPP
2-Hydroxy-ethyl-ThPP
2.3.1.12
1.2.4.1
1.2.4.1
Pyruvate
Acetyl-CoA
S-Acetyldihydro-lipoamide-E
1.8.1.4
Dihydro-lipoamide-E
Lipoamide-E
6.4.1.1
2.3.3.1
2.3.3.3
2.3.3.5
2.3.3.8
Citrate
4.2.1.3
cis-Aconitate
4.2.1.3
Isocitrate
1.1.1.37
Oxaloacetate
(S)-Malate
1.1.5.4
4.2.1.2
Tyrosine metabolism
Arginine biosynthesis
Fumarate
1.1.1.42
Oxalosuccinate
1.1.1.41
1.1.1.286
1.1.1.42
1.3.5.1
1.3.2.4
6.2.1.4
6.2.1.5
2.8.3.18
Oxidative phosphorylation
Succinate
Succinyl-CoA
2.3.1.61
S-Succinyl-dihydrolipoamide-E
1.2.4.2
ThPP
3-Carboxy-1-hydroxypropyl-ThPP
1.2.4.2
2-Oxo-glutarate
Arginine biosynthesis
Ascorbate and aldarate metabolism
Alanine, aspartate and glutamate metabolism
D-Amino acid metabolism
Val , Leu & Ile degradation
Dihydro-lipoamide-E
1.8.1.4
Lipoamide-E
1.2.7.11
1.2.7.3
Data on KEGG graph
Rendered by Pathview

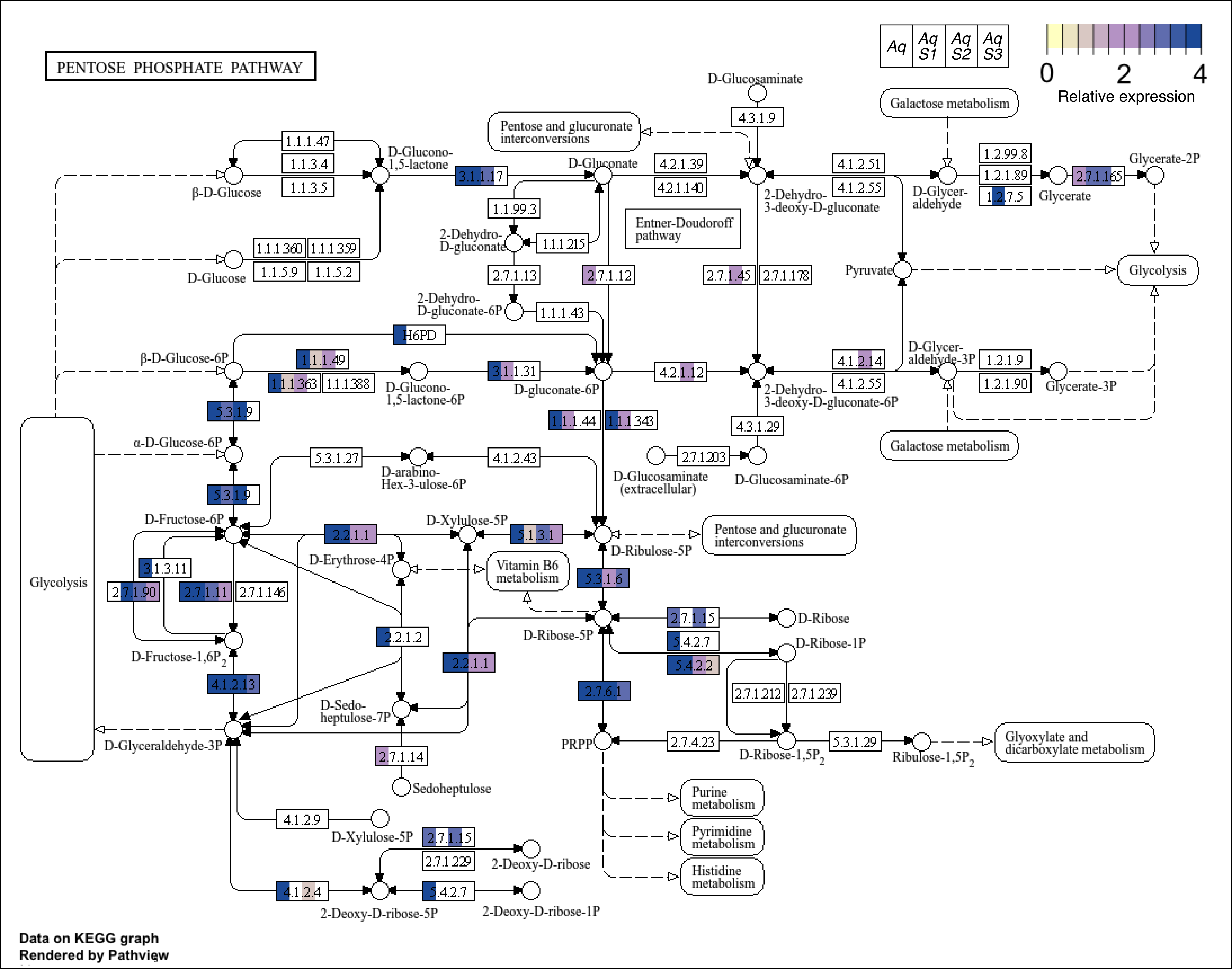
PENTOSE PHOSPHATE PATHWAY
Aq
Aq S1
Aq S2
Aq S3
0
2
4
Relative expression
D-Glucosaminate
4.3.1.9
Galactose metabolism
Pentose and glucuronate interconversions
1.1.1.47
1.1.3.4
1.1.3.5
D-Glucono-1,5-lactone
β-D-Glucose
3.1.1.17
D-Gluconate
4.2.1.39
4.2.1.140
4.1.2.51
4.1.2.55
2-Dehydro-3-deoxy-D-gluconate
D-Glyceraldehyde
1.2.99.8
1.2.1.89
1.2.7.5
Glycerate
2.7.1.165
Glycerate-2P
1.1.99.3
2-Dehydro-D-gluconate
1.1.1.215
Entner-Doudoroff pathway
1.1.1.360
1.1.1.359
D-Glucose
1.1.5.9
1.1.5.2
2.7.1.13
2.7.1.12
2.7.1.45
2.7.1.178
Pyruvate
Glycolysis
2-Dehydro-D-gluconate-6P
1.1.1.43
H6PD
β-D-Glucose-6P
1.1.1.49
1.1.1.363
1.1.1.388
D-Glucono-1,5-lactone-6P
3.1.1.31
D-gluconate-6P
4.2.1.12
4.1.2.14
4.1.2.55
2-Dehydro-3-deoxy-D-gluconate-6P
D-Glyceraldehyde-3P
1.2.1.9
1.2.1.90
Glycerate-3P
5.3.1.9
1.1.1.44
1.1.1.343
4.3.1.29
Galactose metabolism
Glycolysis
α-D-Glucose-6P
5.3.1.27
D-arabino-Hex-3-ulose-6P
4.1.2.43
2.7.1.203
D-Glucosaminate (extracellular)
D-Glucosaminate-6P
5.3.1.9
D-Fructose-6P
2.2.1.1
D-Xylulose-5P
5.1.3.1
D-Ribulose-5P
Pentose and glucuronate interconversions
3.1.3.11
D-Erythrose-4P
Vitamin B6 metabolism
2.7.1.90
2.7.1.11
2.7.1.146
5.3.1.6
2.7.1.15
D-Ribose
D-Ribose-5P
2.2.1.2
5.4.2.7
D-Ribose-1P
D-Fructose-1,6P2
2.2.1.1
5.4.2.2
4.1.2.13
2.7.6.1
2.7.1.212
2.7.1.239
D-Sedo-heptulose-7P
D-Glyceraldehyde-3P
PRPP
2.7.4.23
D-Ribose-1,5P2
5.3.1.29
Ribulose-1,5P2
Glyoxylate and dicarboxylate metabolism
2.7.1.14
Sedoheptulose
Purine metabolism
4.1.2.9
D-Xylulose-5P
2.7.1.15
Pyrimidine metabolism
2.7.1.229
2-Deoxy-D-ribose
Histidine metabolism
4.1.2.4
5.4.2.7
2-Deoxy-D-ribose-5P
2-Deoxy-D-ribose-1P
Data on KEGG graph
Rendered by Pathview

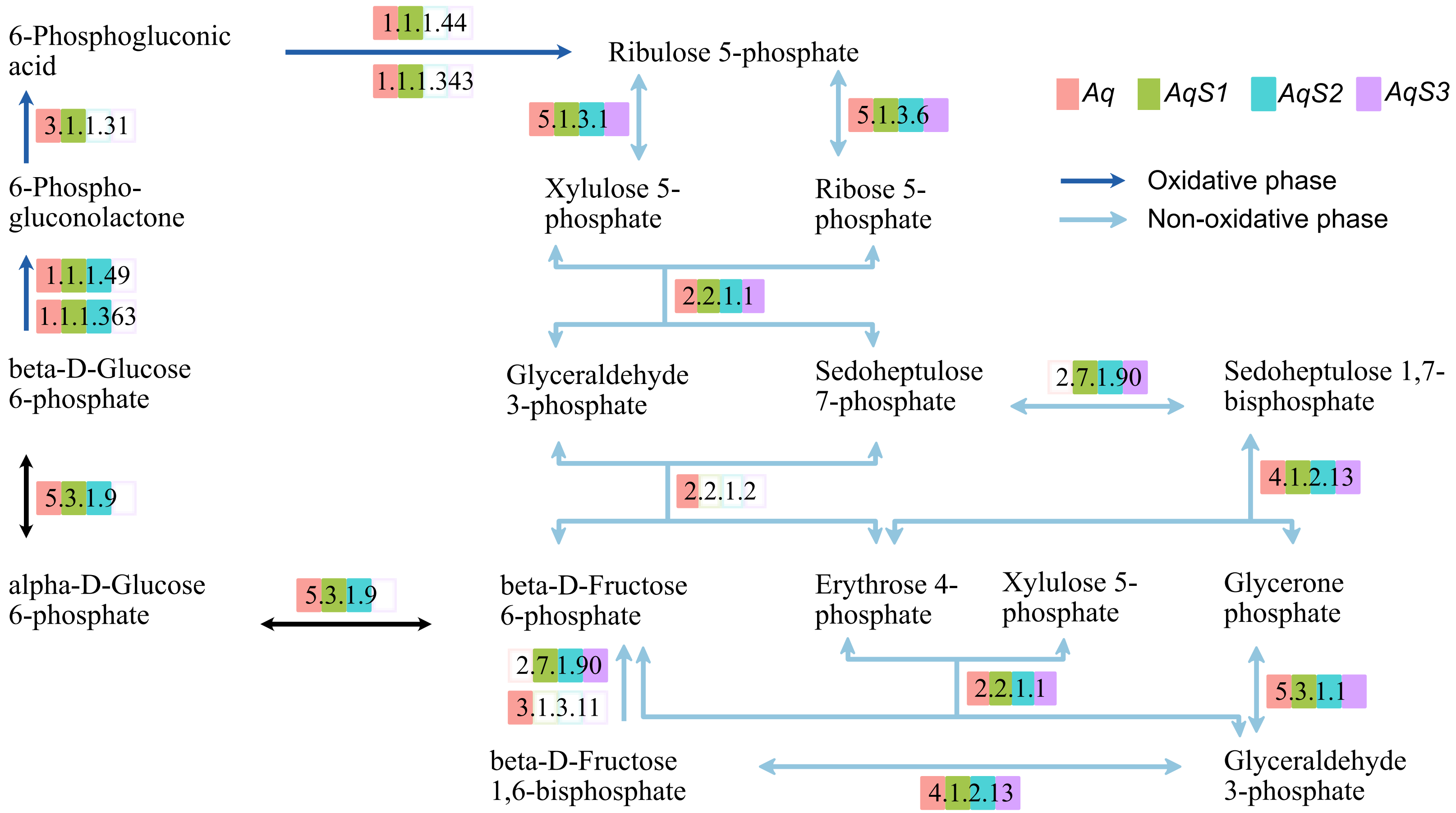
6-Phosphogluconic acid
1.1.1.44
1.1.1.343
Ribulose 5-phosphate
3.1.1.31
5.1.3.1
5.1.3.6
Aq
AqS1
AqS2
AqS3
6-Phospho-gluconolactone
Xylulose 5-phosphate
Ribose 5-phosphate
Oxidative phase
Non-oxidative phase
1.1.1.49
1.1.1.363
2.2.1.1
beta-D-Glucose 6-phosphate
Glyceraldehyde 3-phosphate
Sedoheptulose 7-phosphate
2.7.1.90
Sedoheptulose 1,7-bisphosphate
5.3.1.9
2.2.1.2
4.1.2.13
alpha-D-Glucose 6-phosphate
5.3.1.9
beta-D-Fructose 6-phosphate
Erythrose 4-phosphate
Xylulose 5-phosphate
Glycerone phosphate
2.7.1.90
3.1.3.11
2.2.1.1
5.3.1.1
beta-D-Fructose 1,6-bisphosphate
4.1.2.13
Glyceraldehyde 3-phosphate

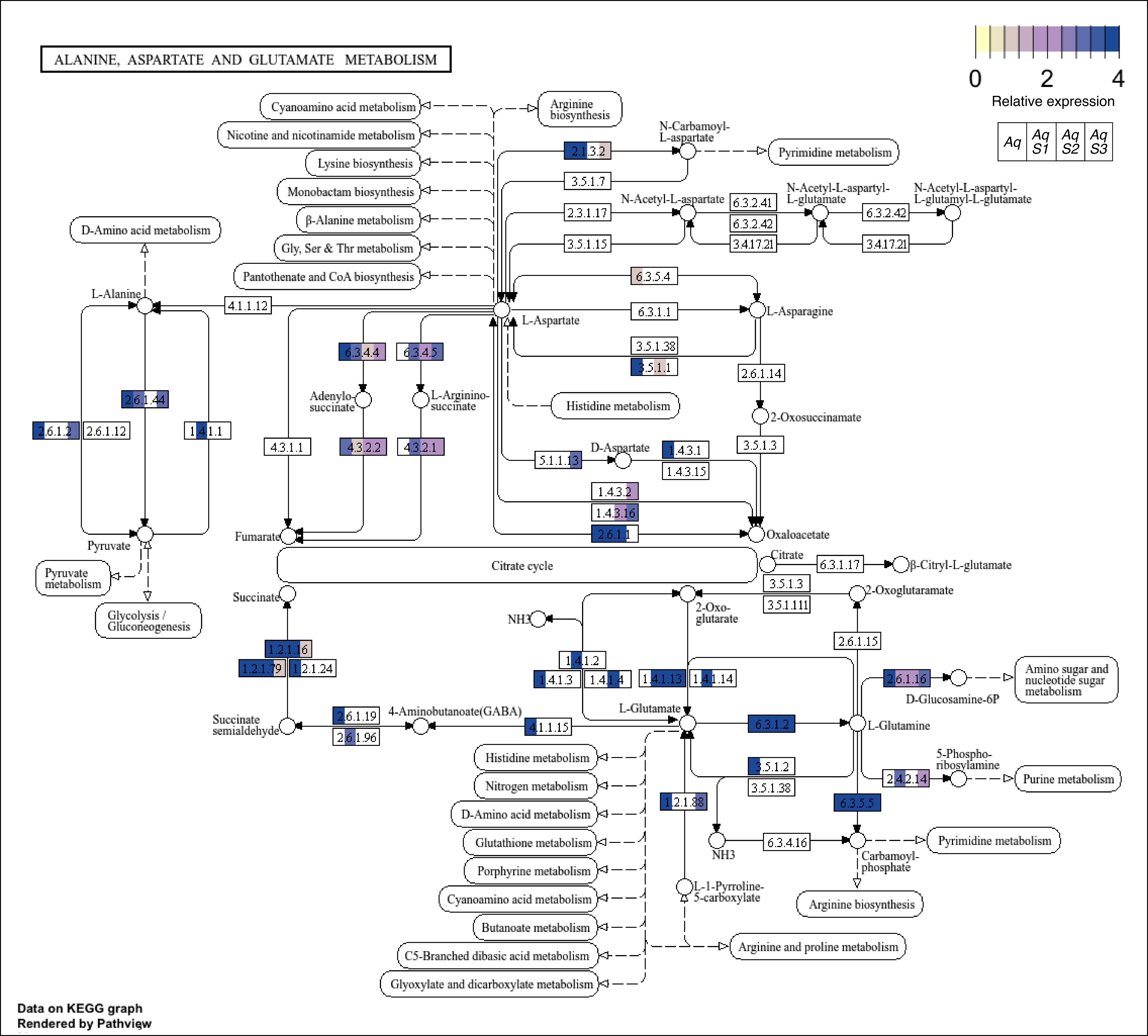

ALANINE, ASPARTATE AND GLUTAMATE METABOLISM
0
2
4
Relative expression
Aq
Aq S1
Aq S2
Aq S3
Cyanoamino acid metabolism
Nicotine and nicotinamide metabolism
Lysine biosynthesis
Monobactam biosynthesis
β-Alanine metabolism
Gly, Ser & Thr metabolism
Pantothenate and CoA biosynthesis
Arginine biosynthesis
D-Amino acid metabolism
N-Carbamoyl-L-aspartate
Pyrimidine metabolism
2.1.3.2
3.5.1.7
2.3.1.17
3.5.1.15
N-Acetyl-L-aspartate
6.3.2.41
6.3.2.42
3.4.17.21
N-Acetyl-L-aspartyl-L-glutamate
N-Acetyl-L-aspartyl-L-glutamyl-L-glutamate
6.3.2.42
3.4.17.21
6.3.5.4
L-Alanine
4.1.1.12
L-Aspartate
6.3.1.1
L-Asparagine
3.5.1.38
3.5.1.1
6.3.4.4
6.3.4.5
Adenylo-succinate
L-Arginino-succinate
2.6.1.44
2.6.1.2
2.6.1.12
1.4.1.1
4.3.1.1
4.3.2.2
4.3.2.1
Histidine metabolism
2.6.1.14
2-Oxosuccinamate
3.5.1.3
D-Aspartate
5.1.1.13
1.4.3.1
1.4.3.15
1.4.3.2
1.4.3.16
2.6.1.1
Oxaloacetate
Pyruvate
Fumarate
Pyruvate metabolism
Glycolysis / Gluconeogenesis
Citrate cycle
Citrate
6.3.1.17
β-Citryl-L-glutamate
3.5.1.3
3.5.1.111
Succinate
2-Oxoglutaramate
NH3
2-Oxo-glutarate
1.2.1.16
1.2.1.79
1.2.1.24
1.4.1.2
1.4.1.3
1.4.1.4
1.4.1.13
1.4.1.14
2.6.1.15
2.6.1.16
D-Glucosamine-6P
Amino sugar and nucleotide sugar metabolism
Succinate semialdehyde
2.6.1.19
2.6.1.96
4-Aminobutanoate(GABA)
4.1.1.15
L-Glutamate
6.3.1.2
L-Glutamine
5-Phospho-ribosylamine
2.4.2.14
Purine metabolism
Histidine metabolism
Nitrogen metabolism
D-Amino acid metabolism
Glutathione metabolism
Porphyrine metabolism
Cyanoamino acid metabolism
Butanoate metabolism
C5-Branched dibasic acid metabolism
Glyoxylate and dicarboxylate metabolism
3.5.1.2
3.5.1.38
1.2.1.88
6.3.5.5
NH3
6.3.4.16
Carbamoyl-phosphate
Pyrimidine metabolism
Arginine biosynthesis
L-1-Pyrroline-5-carboxylate
Arginine and proline metabolism
Data on KEGG graph
Rendered by Pathview

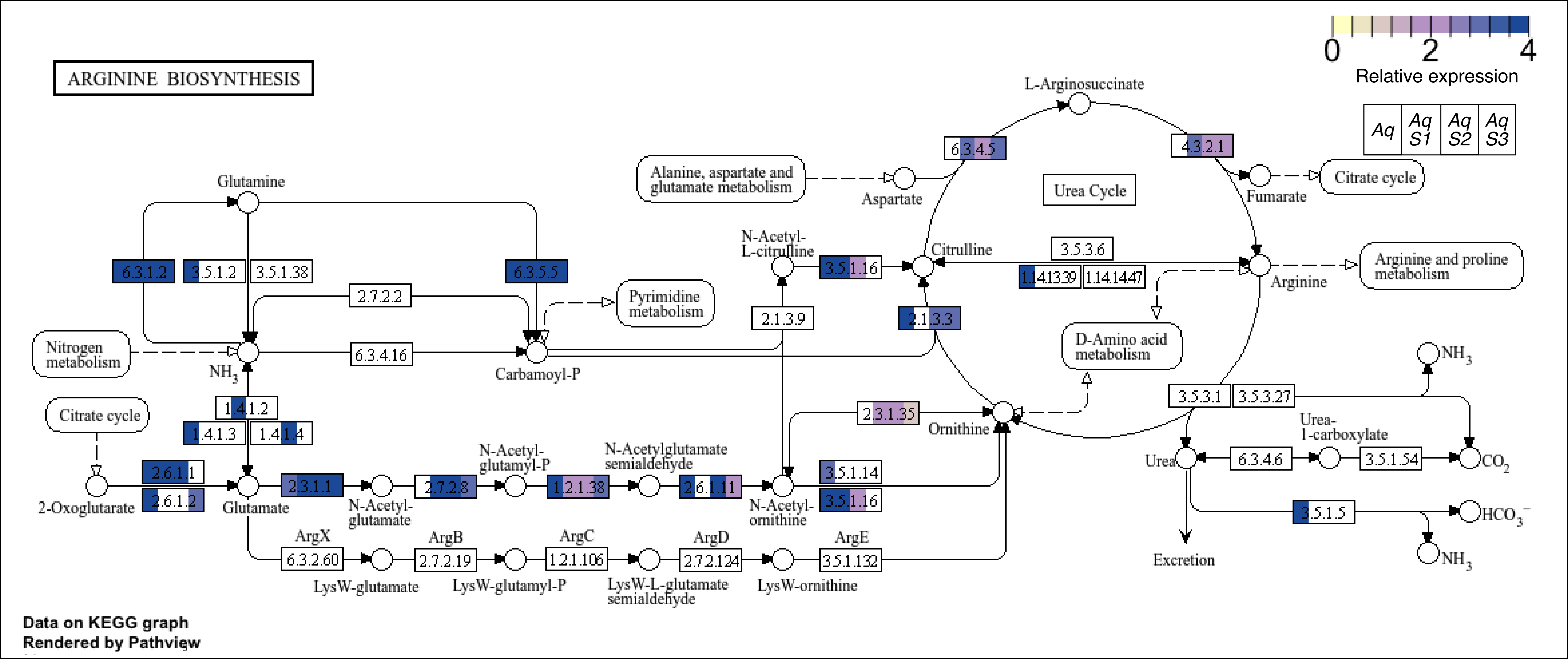

ARGININE BIOSYNTHESIS
0
2
4
Relative expression
Aq
Aq S1
Aq S2
Aq S3
L-Arginosuccinate
6.3.4.5
4.3.2.1
Alanine, aspartate and glutamate metabolism
Aspartate
Urea Cycle
Fumarate
Citrate cycle
Glutamine
N-Acetyl-L-citrulline
Citrulline
3.5.3.6
Arginine
Arginine and proline metabolism
6.3.1.2
3.5.1.2
3.5.1.38
6.3.5.5
3.5.1.16
1.14.13.39
1.14.14.47
2.7.2.2
Pyrimidine metabolism
2.1.3.9
2.1.3.3
D-Amino acid metabolism
Nitrogen metabolism
NH3
6.3.4.16
Carbamoyl-P
NH3
1.4.1.2
3.5.3.1
3.5.3.27
Citrate cycle
1.4.1.3
1.4.1.4
2.3.1.35
Ornithine
Urea-1-carboxylate
2.6.1.1
N-Acetyl-glutamyl-P
N-Acetylglutamate semialdehyde
Urea
6.3.4.6
3.5.1.54
CO2
2-Oxoglutarate
2.6.1.2
Glutamate
2.3.1.1
N-Acetyl-glutamate
2.7.2.8
1.2.1.38
2.6.1.11
N-Acetyl-ornithine
3.5.1.14
3.5.1.16
3.5.1.5
HCO3−
ArgX
ArgB
ArgC
ArgD
ArgE
Excretion
6.3.2.60
2.7.2.19
1.2.1.106
2.7.2.124
3.5.1.132
NH3
LysW-glutamate
LysW-glutamyl-P
LysW-L-glutamate semialdehyde
LysW-ornithine
Data on KEGG graph
Rendered by Pathview

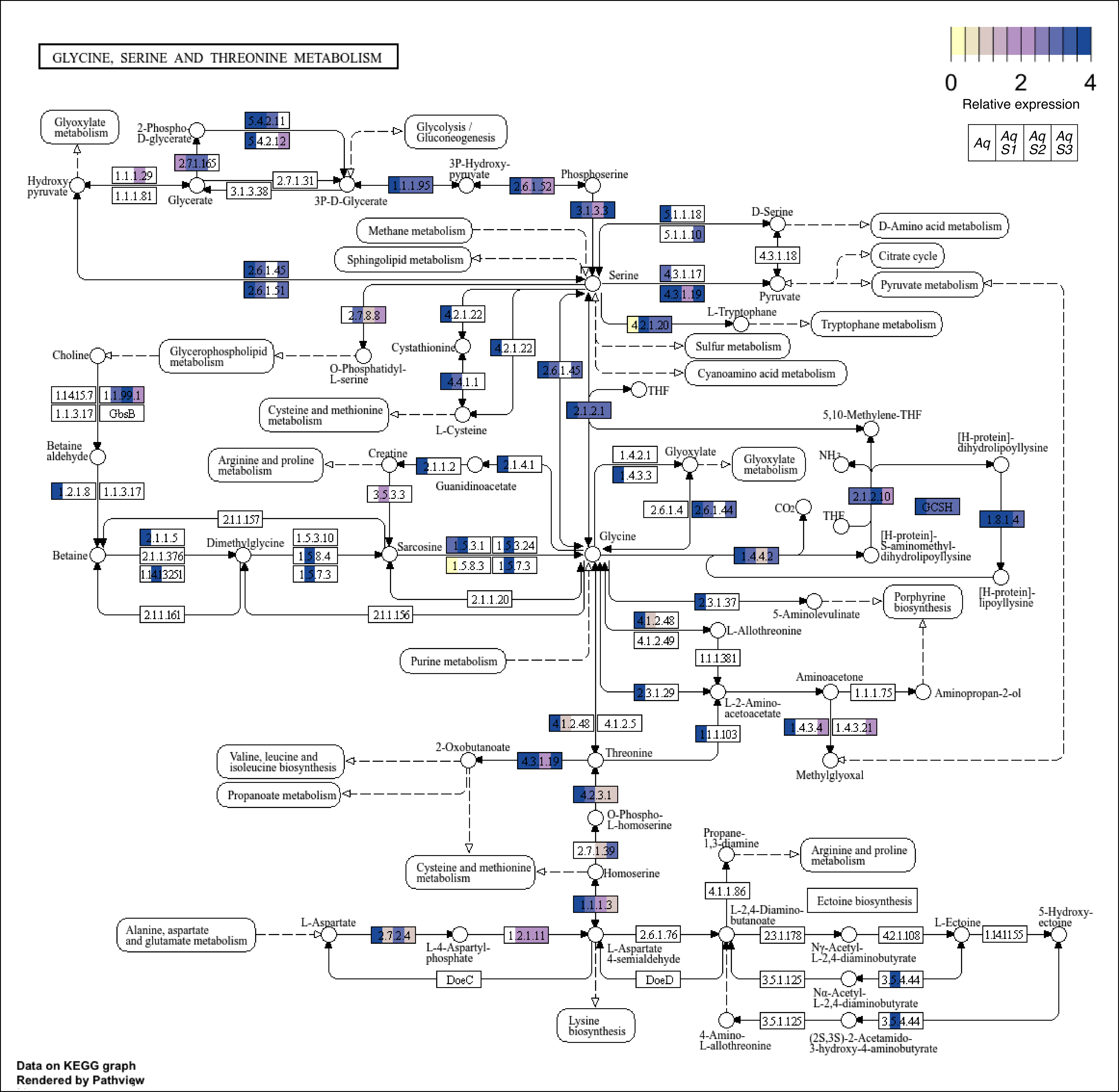
GLYCINE, SERINE AND THREONINE METABOLISM
0
2
4
Relative expression
Aq
Aq S1
Aq S2
Aq S3
Glyoxylate metabolism
2-Phospho-D-glycerate
5.4.2.11
5.4.2.12
Glycolysis / Gluconeogenesis
2.7.1.165
3P-Hydroxy-pyruvate
Phosphoserine
Hydroxy-pyruvate
1.1.1.29
1.1.1.81
Glycerate
2.7.1.31
3.1.3.38
3P-D-Glycerate
1.1.1.95
2.6.1.52
3.1.3.3
5.1.1.18
5.1.1.10
D-Serine
D-Amino acid metabolism
Methane metabolism
Sphingolipid metabolism
4.3.1.18
Citrate cycle
Pyruvate metabolism
2.6.1.45
2.6.1.51
Serine
4.3.1.17
4.3.1.19
Pyruvate
2.7.8.8
4.2.1.22
L-Tryptophane
4.2.1.20
Tryptophane metabolism
Sulfur metabolism
Choline
Glycerophospholipid metabolism
O-Phosphatidyl-L-serine
Cystathionine
4.2.1.22
2.6.1.45
Cyanoamino acid metabolism
1.1.4.15.7
1.1.99.1
1.1.3.17
GbsB
4.4.1.1
THF
Cysteine and methionine metabolism
L-Cysteine
2.1.2.1
5,10-Methylene-THF
Betaine aldehyde
[H-protein]-dihydrolipoyllysine
Arginine and proline metabolism
Creatine
2.1.1.2
2.1.4.1
1.4.2.1
Glyoxylate
Glyoxylate metabolism
$NH_3$
1.2.1.8
1.1.3.17
3.5.3.3
Guanidinoacetate
1.4.3.3
2.1.2.10
GCSH
2.6.1.4
2.6.1.44
$CO_2$
THF
1.8.1.4
2.1.1.157
Betaine
2.1.1.5
2.1.1.376
1.1.4.13251
Dimethylglycine
1.5.3.10
1.5.8.4
1.5.7.3
Sarcosine
1.5.3.1
1.5.3.24
1.5.8.3
1.5.7.3
Glycine
1.4.4.2
[H-protein]-S-aminomethyl-dihydrolipoyllysine
[H-protein]-lipoyllysine
2.1.1.20
2.1.1.161
2.1.1.156
2.3.1.37
5-Aminolevulinate
Porphyrine biosynthesis
4.1.2.48
4.1.2.49
L-Allothreonine
Purine metabolism
1.1.1.381
Aminoacetone
2.3.1.29
1.1.1.75
Aminopropan-2-ol
L-2-Amino-acetoacetate
4.1.2.48
4.1.2.5
1.1.1.103
1.4.3.4
1.4.3.21
2-Oxobutanoate
Valine, leucine and isoleucine biosynthesis
4.3.1.19
Threonine
Methylglyoxal
Propanoate metabolism
4.2.3.1
O-Phospho-L-homoserine
2.7.1.39
Propane-1,3-diamine
Arginine and proline metabolism
Cysteine and methionine metabolism
Homoserine
4.1.1.86
Ectoine biosynthesis
1.1.1.3
L-2,4-Diamino-butanoate
L-Aspartate
L-Ectoine
5-Hydroxy-ectoine
Alanine, aspartate and glutamate metabolism
2.7.2.4
L-4-Aspartyl-phosphate
1.2.1.11
2.6.1.76
L-Aspartate 4-semialdehyde
2.3.1.178
Nγ-Acetyl-L-2,4-diaminobutyrate
4.2.1.108
1.14.11.55
DoeC
DoeD
3.5.1.125
3.5.4.44
Nα-Acetyl-L-2,4-diaminobutyrate
Lysine biosynthesis
4-Amino-L-allothreonine
3.5.1.125
3.5.4.44
(2S,3S)-2-Acetamido-3-hydroxy-4-aminobutyrate
Data on KEGG graph
Rendered by Pathview

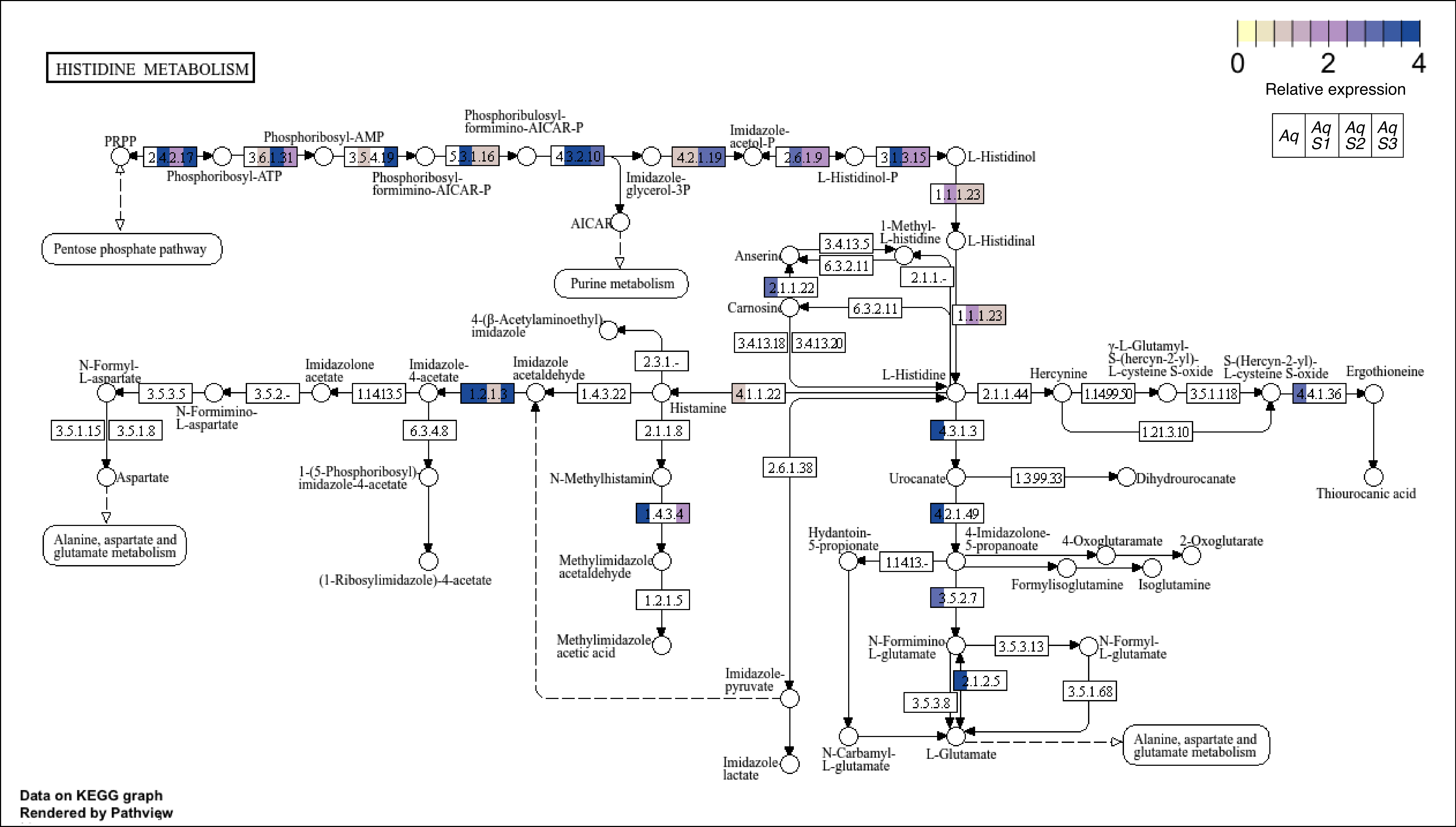
HISTIDINE METABOLISM
0
2
4
Relative expression
Aq
Aq S1
Aq S2
Aq S3
PRPP
2.4.2.17
Phosphoribosyl-ATP
3.6.1.31
Phosphoribosyl-AMP
3.5.4.19
Phosphoribosyl-formimino-AICAR-P
5.3.1.16
Phosphoribulosyl-formimino-AICAR-P
4.3.2.10
Imidazole-glycerol-3P
AICAR
Purine metabolism
4.2.1.19
Imidazole-acetol-P
2.6.1.9
L-Histidinol-P
3.1.3.15
L-Histidinol
1.1.1.23
L-Histidinal
Pentose phosphate pathway
1-Methyl-L-histidine
3.4.13.5
6.3.2.11
Anserine
2.1.1.-
2.1.1.22
Carnosine
6.3.2.11
1.1.1.23
3.4.13.18
3.4.13.20
4-(β-Acetylaminoethyl)imidazole
2.3.1.-
N-Formyl-L-aspartate
3.5.3.5
N-Formimino-L-aspartate
3.5.2.-
Imidazolone acetate
1.14.13.5
Imidazole-4-acetate
1.2.1.3
Imidazole acetaldehyde
1.4.3.22
Histamine
4.1.1.22
L-Histidine
2.1.1.44
Hercynine
1.14.99.50
γ-L-Glutamyl-S-(hercyn-2-yl)-L-cysteine S-oxide
3.5.1.118
S-(Hercyn-2-yl)-L-cysteine S-oxide
4.4.1.36
Ergothioneine
1.21.3.10
Thiourocanic acid
3.5.1.15
3.5.1.8
Aspartate
Alanine, aspartate and glutamate metabolism
6.3.4.8
1-(5-Phosphoribosyl)imidazole-4-acetate
(1-Ribosylimidazole)-4-acetate
2.1.1.8
N-Methylhistamine
1.4.3.4
Methylimidazole acetaldehyde
1.2.1.5
Methylimidazole acetic acid
2.6.1.38
Imidazole-pyruvate
Imidazole lactate
4.3.1.3
Urocanate
1.3.99.33
Dihydrourocanate
4.2.1.49
4-Imidazolone-5-propanoate
Hydantoin-5-propionate
1.14.13.-
4-Oxoglutaramate
2-Oxoglutarate
Formylisoglutamine
Isoglutamine
3.5.2.7
N-Formimino-L-glutamate
3.5.3.13
N-Formyl-L-glutamate
2.1.2.5
3.5.3.8
3.5.1.68
N-Carbamyl-L-glutamate
L-Glutamate
Alanine, aspartate and glutamate metabolism
Data on KEGG graph
Rendered by Pathview

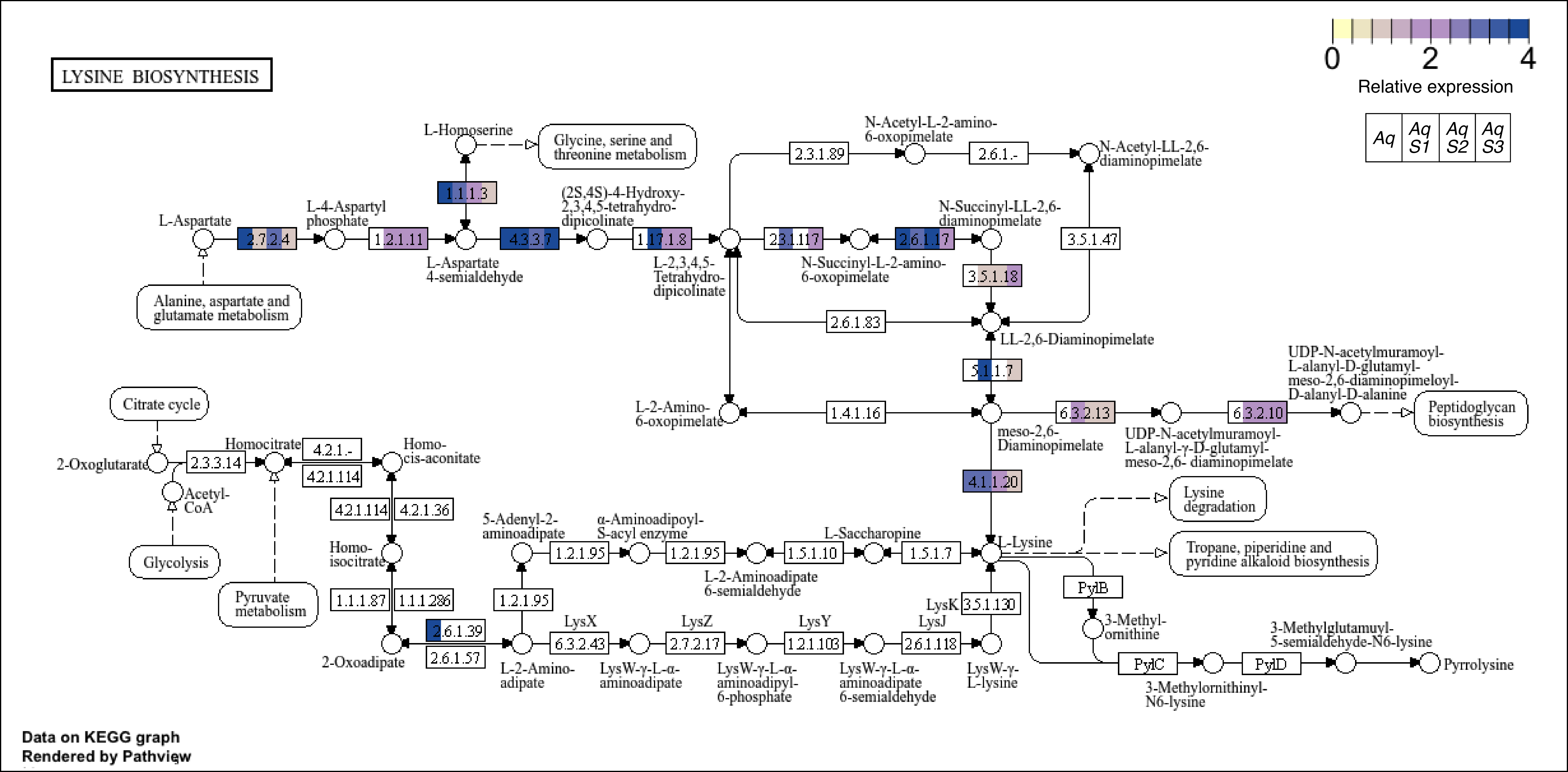

LYSINE BIOSYNTHESIS
0
2
4
Relative expression
Aq
Aq S1
Aq S2
Aq S3
L-Aspartate
2.7.2.4
L-4-Aspartyl phosphate
1.2.1.11
L-Aspartate 4-semialdehyde
L-Homoserine
1.1.1.3
Glycine, serine and threonine metabolism
4.3.3.7
(2S,4S)-4-Hydroxy-2,3,4,5-tetrahydro-dipicolinate
1.17.1.8
L-2,3,4,5-Tetrahydro-dipicolinate
Alanine, aspartate and glutamate metabolism
2.3.1.89
N-Acetyl-L-2-amino-6-oxopimelate
2.6.1.-
N-Acetyl-LL-2,6-diaminopimelate
2.3.1.117
N-Succinyl-L-2-amino-6-oxopimelate
2.6.1.17
N-Succinyl-LL-2,6-diaminopimelate
3.5.1.47
3.5.1.18
2.6.1.83
LL-2,6-Diaminopimelate
5.1.1.7
L-2-Amino-6-oxopimelate
1.4.1.16
meso-2,6-Diaminopimelate
6.3.2.13
UDP-N-acetylmuramoyl-L-alanyl-γ-D-glutamyl-meso-2,6- diaminopimelate
6.3.2.10
UDP-N-acetylmuramoyl-L-alanyl-D-glutamyl-meso-2,6-diaminopimeloyl-D-alanyl-D-alanine
Peptidoglycan biosynthesis
4.1.1.20
L-Lysine
Lysine degradation
Tropane, piperidine and pyridine alkaloid biosynthesis
Citrate cycle
2-Oxoglutarate
2.3.3.14
Homocitrate
4.2.1.-
4.2.1.114
Homo-cis-aconitate
Acetyl-CoA
Glycolysis
Pyruvate metabolism
4.2.1.114
4.2.1.36
Homo-isocitrate
1.1.1.87
1.1.1.286
2-Oxoadipate
2.6.1.39
2.6.1.57
L-2-Amino-adipate
1.2.1.95
5-Adenyl-2-aminoadipate
1.2.1.95
α-Aminoadipoyl-S-acyl enzyme
1.2.1.95
L-2-Aminoadipate 6-semialdehyde
1.5.1.10
L-Saccharopine
1.5.1.7
LysX
6.3.2.43
LysW-γ-L-α-aminoadipate
LysZ
2.7.2.17
LysW-γ-L-α-aminoadipyl-6-phosphate
LysY
1.2.1.103
LysW-γ-L-α-aminoadipate 6-semialdehyde
LysJ
2.6.1.118
LysW-γ-L-lysine
LysK
3.5.1.130
PylB
3-Methyl-ornithine
PylC
3-Methylornithinyl-N6-lysine
PylD
3-Methylglutamuyl-5-semialdehyde-N6-lysine
Pyrrolysine
Data on KEGG graph
Rendered by Pathview

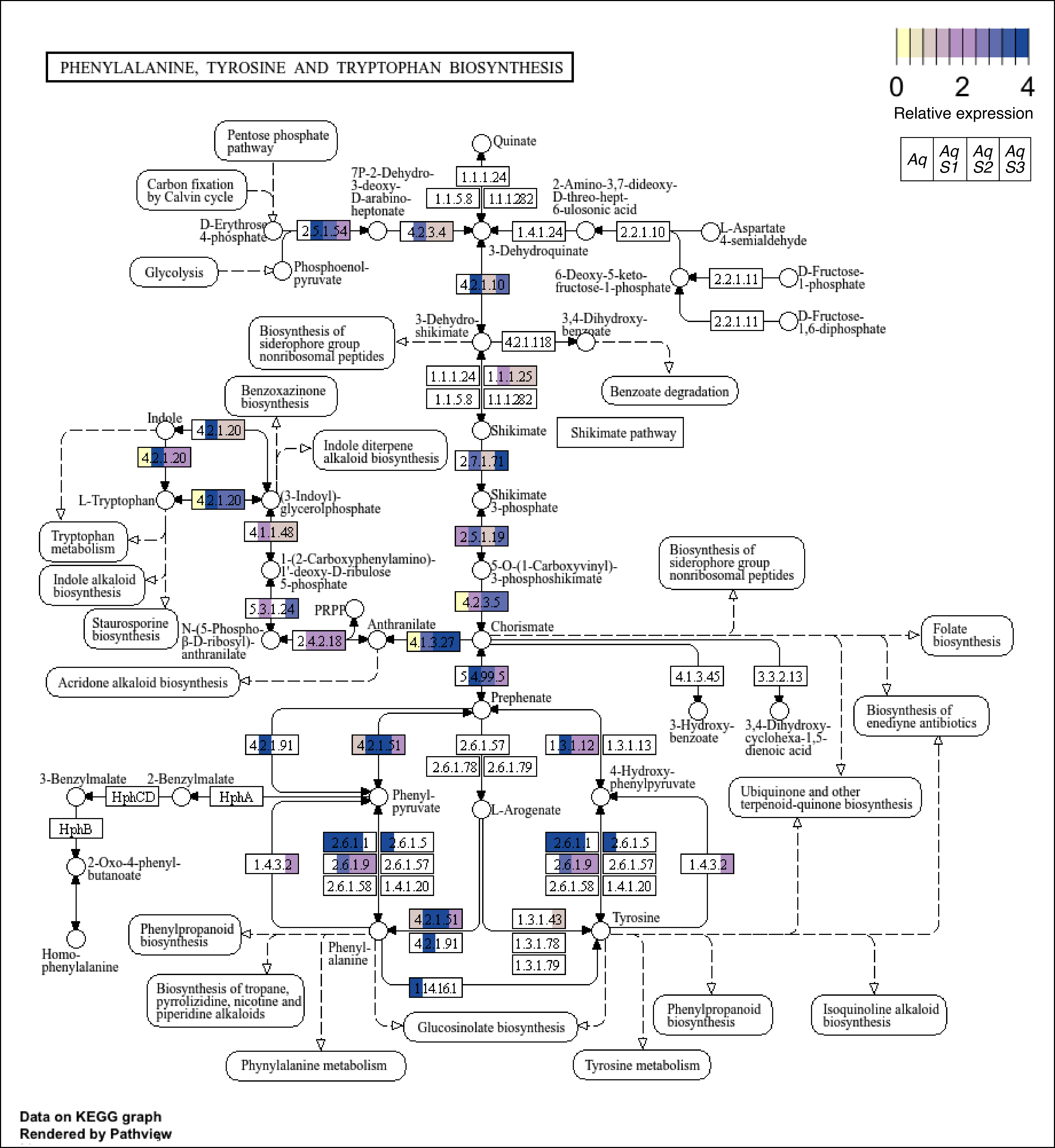
PHENYLALANINE, TYROSINE AND TRYPTOPHAN BIOSYNTHESIS
0
2
4
Relative expression
Aq
Aq S1
Aq S2
Aq S3
Pentose phosphate pathway
Carbon fixation by Calvin cycle
D-Erythrose 4-phosphate
Glycolysis
Phosphoenol-pyruvate
2.5.1.54
7P-2-Dehydro-3-deoxy-D-arabino-heptonate
4.2.3.4
Quinate
1.1.1.24
1.1.5.8
1.1.1.282
2-Amino-3,7-dideoxy-D-threo-hept-6-ulosonic acid
1.4.1.24
2.2.1.10
L-Aspartate 4-semialdehyde
3-Dehydroquinate
4.2.1.10
6-Deoxy-5-keto-fructose-1-phosphate
2.2.1.11
D-Fructose-1-phosphate
2.2.1.11
D-Fructose-1,6-diphosphate
Biosynthesis of siderophore group nonribosomal peptides
3-Dehydro-shikimate
4.2.1.118
3,4-Dihydroxy-benzoate
1.1.1.24
1.1.1.25
1.1.5.8
1.1.1.282
Benzoate degradation
Benzoxazinone biosynthesis
Indole
4.2.1.20
Shikimate
Shikimate pathway
Indole diterpene alkaloid biosynthesis
4.2.1.20
2.7.1.71
L-Tryptophan
4.2.1.20
(3-Indoyl)-glycerolphosphate
Shikimate 3-phosphate
Tryptophan metabolism
4.1.1.48
2.5.1.19
Biosynthesis of siderophore group nonribosomal peptides
Indole alkaloid biosynthesis
1-(2-Carboxyphenylamino)-1'-deoxy-D-ribulose 5-phosphate
5-O-(1-Carboxyvinyl)-3-phosphoshikimate
5.3.1.24
4.2.3.5
PRPP
Staurosporine biosynthesis
N-(5-Phospho-β-D-ribosyl)-anthranilate
2.4.2.18
Anthranilate
4.1.3.27
Chorismate
Folate biosynthesis
Acridone alkaloid biosynthesis
5.4.99.5
4.1.3.45
3.3.2.13
Prephenate
3-Hydroxy-benzoate
3,4-Dihydroxy-cyclohexa-1,5-dienoic acid
Biosynthesis of enediyne antibiotics
4.2.1.91
4.2.1.51
2.6.1.57
1.3.1.12
1.3.1.13
2.6.1.78
2.6.1.79
4-Hydroxy-phenylpyruvate
3-Benzylmalate
2-Benzylmalate
HphCD
HphA
Phenyl-pyruvate
L-Arogenate
Ubiquinone and other terpenoid-quinone biosynthesis
HphB
2.6.1.1
2.6.1.5
2.6.1.1
2.6.1.5
1.4.3.2
2.6.1.9
2.6.1.57
2.6.1.9
2.6.1.57
1.4.3.2
2-Oxo-4-phenyl-butanoate
2.6.1.58
1.4.1.20
2.6.1.58
1.4.1.20
4.2.1.51
1.3.1.43
Tyrosine
Phenylpropanoid biosynthesis
4.2.1.91
1.3.1.78
Phenyl-alanine
Homo-phenylalanine
1.3.1.79
1.14.16.1
Biosynthesis of tropane, pyrrolizidine, nicotine and piperidine alkaloids
Glucosinolate biosynthesis
Phenylpropanoid biosynthesis
Isoquinoline alkaloid biosynthesis
Phynylalanine metabolism
Tyrosine metabolism
Data on KEGG graph
Rendered by Pathview

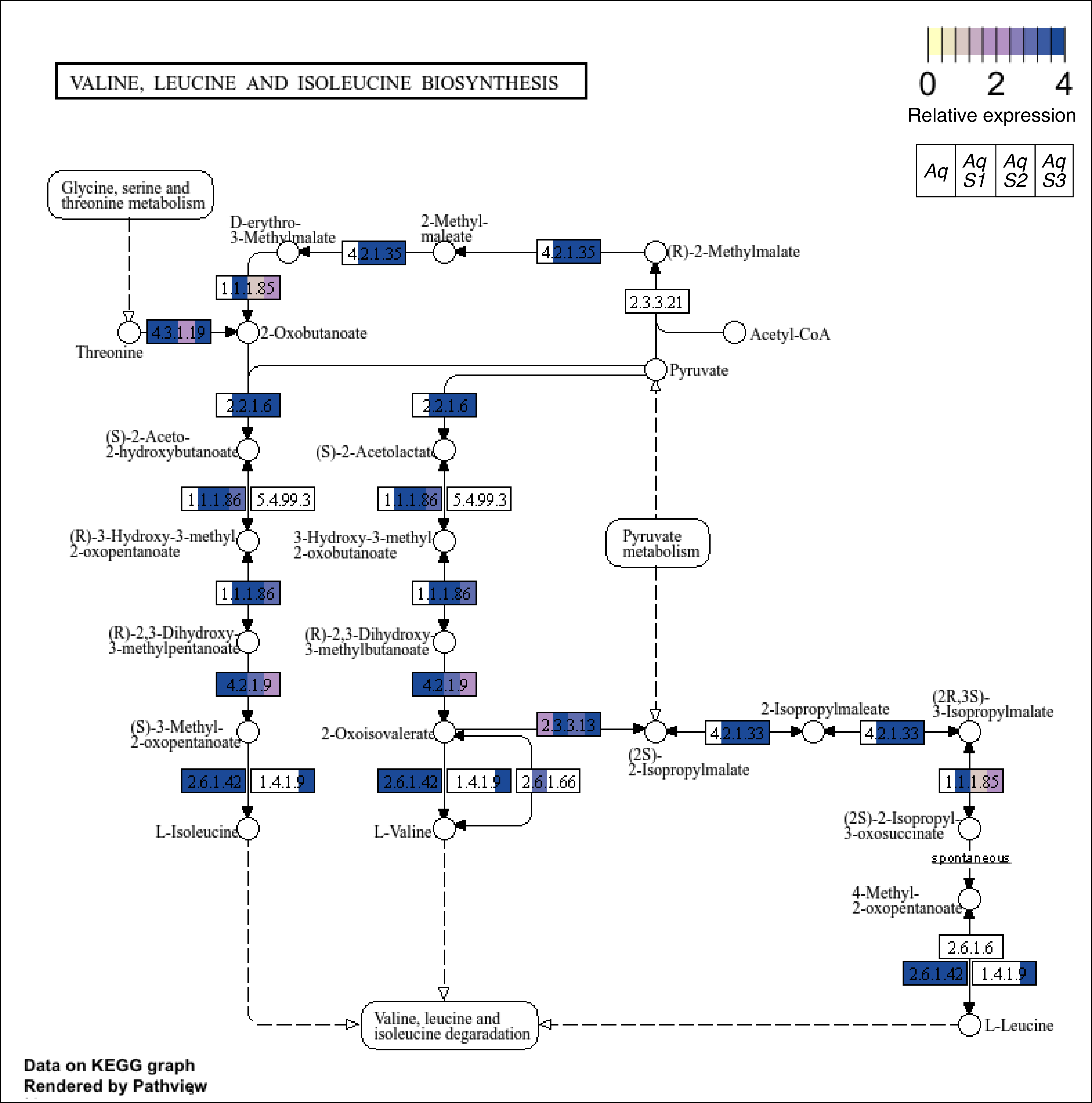
VALINE, LEUCINE AND ISOLEUCINE BIOSYNTHESIS
0
2
4
Relative expression
Aq
Aq S1
Aq S2
Aq S3
Glycine, serine and threonine metabolism
D-erythro-3-Methylmalate
2-Methyl-maleate
4.2.1.35
4.2.1.35
(R)-2-Methylmalate
1.1.1.85
2.3.3.21
Threonine
4.3.1.19
2-Oxobutanoate
Acetyl-CoA
Pyruvate
2.2.1.6
2.2.1.6
(S)-2-Aceto-2-hydroxybutanoate
(S)-2-Acetolactate
1.1.1.86
5.4.99.3
1.1.1.86
5.4.99.3
(R)-3-Hydroxy-3-methyl-2-oxopentanoate
3-Hydroxy-3-methyl-2-oxobutanoate
Pyruvate metabolism
1.1.1.86
1.1.1.86
(R)-2,3-Dihydroxy-3-methylpentanoate
(R)-2,3-Dihydroxy-3-methylbutanoate
4.2.1.9
4.2.1.9
(S)-3-Methyl-2-oxopentanoate
2-Oxoisovalerate
2.3.3.13
(2S)-2-Isopropylmalate
4.2.1.33
2-Isopropylmaleate
4.2.1.33
(2R,3S)-3-Isopropylmalate
2.6.1.42
1.4.1.9
2.6.1.42
1.4.1.9
2.6.1.66
1.1.1.85
L-Isoleucine
L-Valine
(2S)-2-Isopropyl-3-oxosuccinate
spontaneous
4-Methyl-2-oxopentanoate
2.6.1.6
2.6.1.42
1.4.1.9
Valine, leucine and isoleucine degaradation
L-Leucine
Data on KEGG graph
Rendered by Pathview

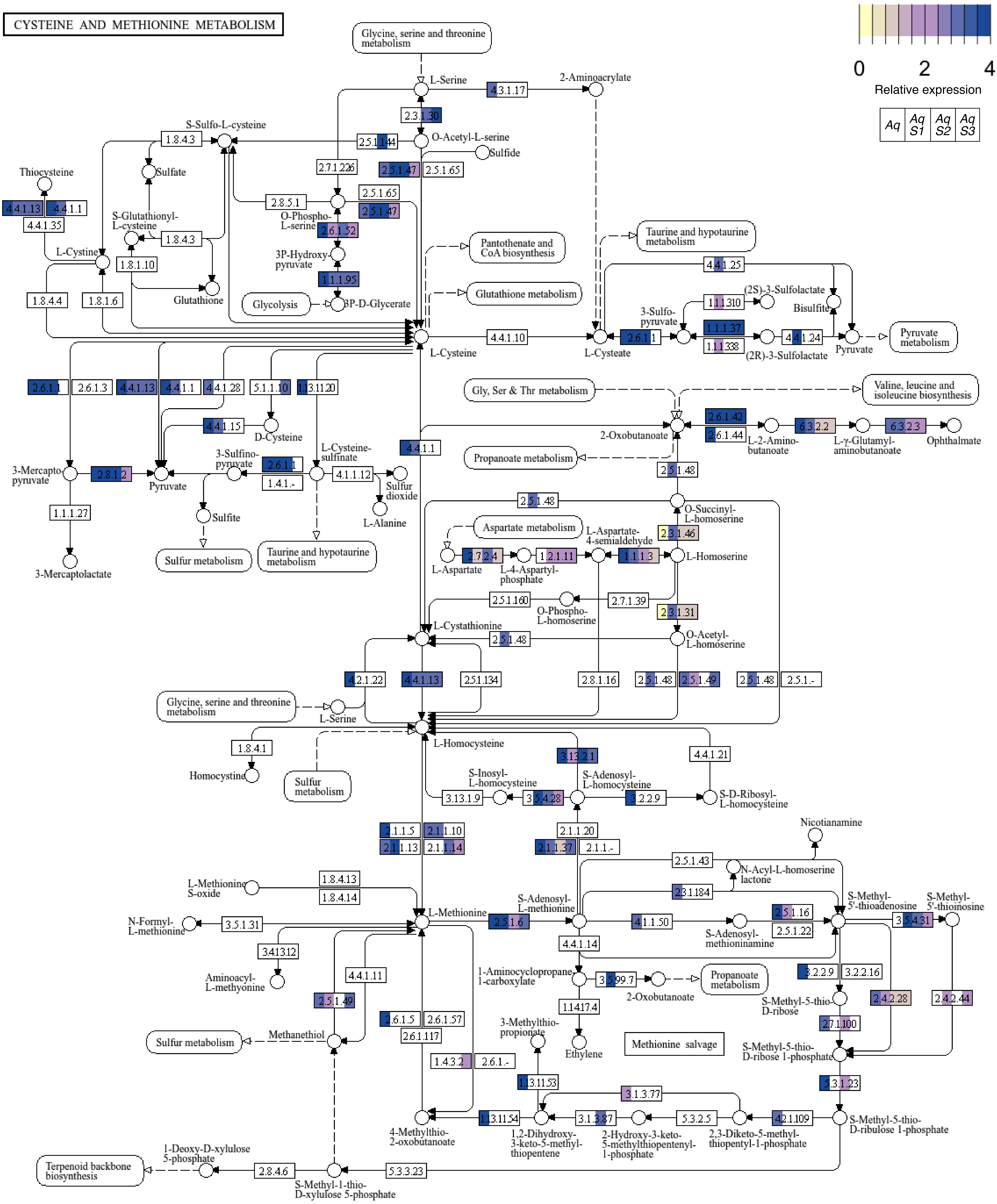


**Data on KEGG graph**
**Rendered by Pathview**

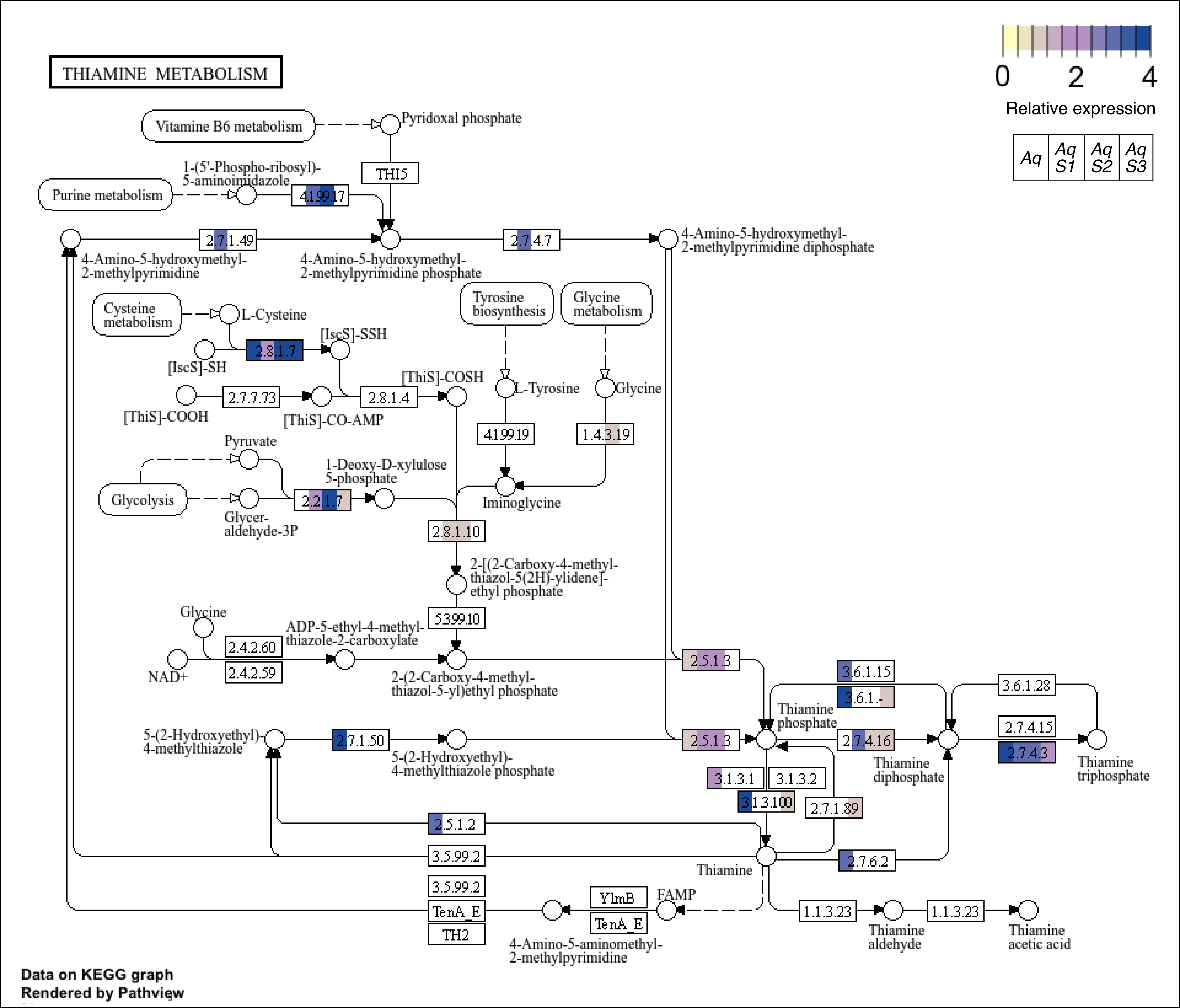
THIAMINE METABOLISM
0
2
4
Relative expression
Aq
Aq S1
Aq S2
Aq S3
Vitamine B6 metabolism
Pyridoxal phosphate
THI5
Purine metabolism
1-(5'-Phospho-ribosyl)-
5-aminoimidazole
4.1.99.17
2.7.1.49
2.7.4.7
4-Amino-5-hydroxymethyl-
2-methylpyrimidine diphosphate
4-Amino-5-hydroxymethyl-
2-methylpyrimidine
4-Amino-5-hydroxymethyl-
2-methylpyrimidine phosphate
Cysteine metabolism
L-Cysteine
[IscS]-SSH
2.8.1.7
[IscS]-SH
Tyrosine biosynthesis
Glycine metabolism
[ThiS]-COSH
L-Tyrosine
Glycine
2.7.7.73
2.8.1.4
[ThiS]-COOH
[ThiS]-CO-AMP
4.1.99.19
1.4.3.19
Pyruvate
1-Deoxy-D-xylulose
5-phosphate
Glycolysis
2.2.1.7
Iminoglycine
Glycer-
aldehyde-3P
2.8.1.10
2-[(2-Carboxy-4-methyl-
thiazol-5(2H)-ylidene]-
ethyl phosphate
5.3.99.10
Glycine
ADP-5-ethyl-4-methyl-
thiazole-2-carboxylate
2.4.2.60
2.4.2.59
NAD+
2-(2-Carboxy-4-methyl-
thiazol-5-yl)ethyl phosphate
2.5.1.3
3.6.1.15
3.6.1.-
3.6.1.28
Thiamine
phosphate
5-(2-Hydroxyethyl)-
4-methylthiazole
2.7.1.50
5-(2-Hydroxyethyl)-
4-methylthiazole phosphate
2.5.1.3
2.7.4.16
2.7.4.15
2.7.4.3
Thiamine
diphosphate
Thiamine
triphosphate
3.1.3.1
3.1.3.2
3.1.3.100
2.7.1.89
2.5.1.2
3.5.99.2
Thiamine
2.7.6.2
3.5.99.2
TenA_E
TH2
YlmB
TenA_E
FAMP
1.1.3.23
1.1.3.23
Thiamine
aldehyde
Thiamine
acetic acid
4-Amino-5-aminomethyl-
2-methylpyrimidine
Data on KEGG graph
Rendered by Pathview

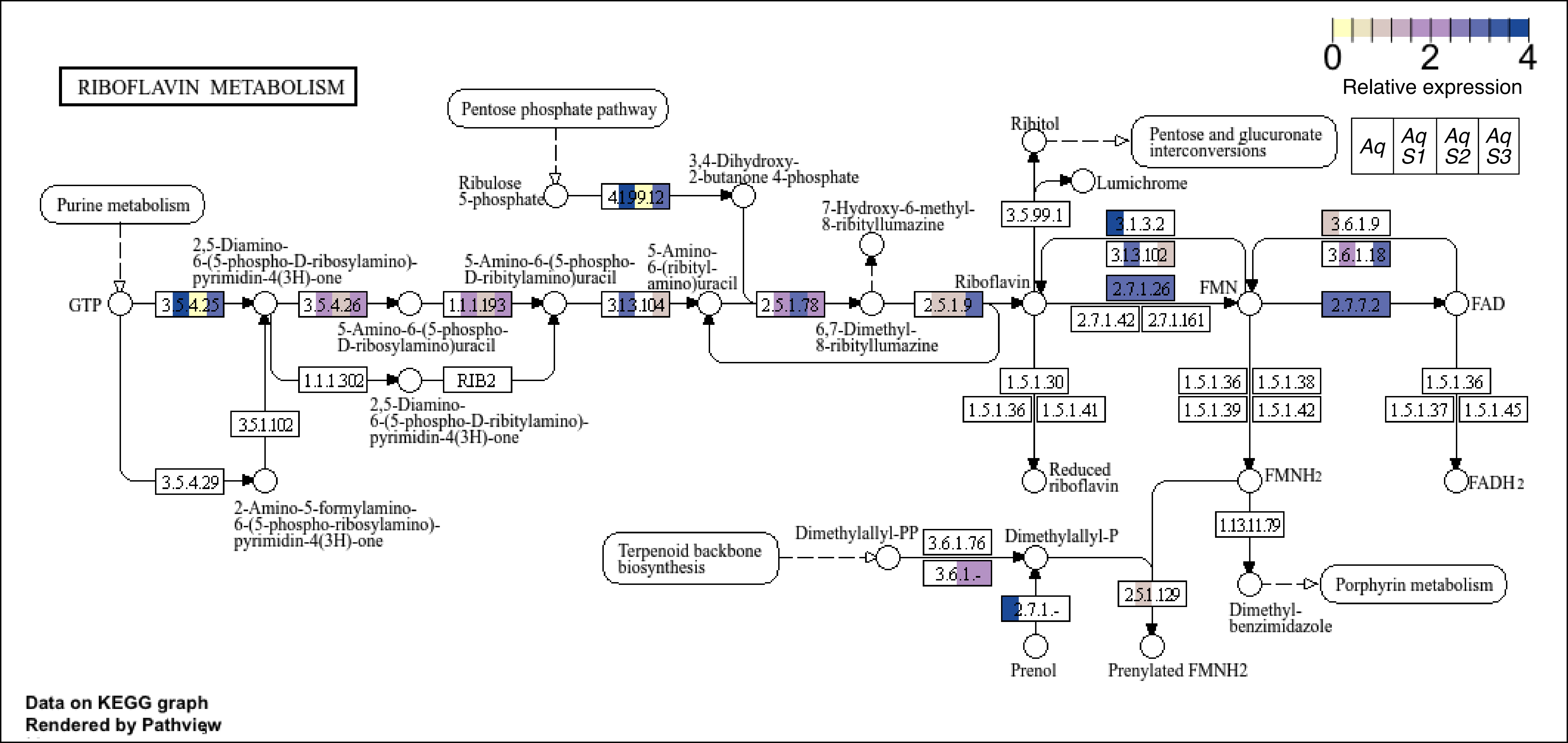
RIBOFLAVIN METABOLISM
0
2
4
Relative expression
Aq
Aq S1
Aq S2
Aq S3
Pentose phosphate pathway
Purine metabolism
Ribulose 5-phosphate
4.1.99.12
3,4-Dihydroxy-2-butanone 4-phosphate
Ribitol
Pentose and glucuronate interconversions
Lumichrome
3.5.99.1
7-Hydroxy-6-methyl-8-ribityllumazine
3.1.3.2
3.1.3.102
3.6.1.9
3.6.1.18
2.7.1.26
2.7.7.2
2.7.1.42
2.7.1.161
GTP
3.5.4.25
2,5-Diamino-6-(5-phospho-D-ribosylamino)-pyrimidin-4(3H)-one
3.5.4.26
5-Amino-6-(5-phospho-D-ribosylamino)uracil
1.1.1.193
5-Amino-6-(5-phospho-D-ribitylamino)uracil
3.1.3.104
5-Amino-6-(ribityl-amino)uracil
2.5.1.78
6,7-Dimethyl-8-ribityllumazine
2.5.1.9
Riboflavin
FMN
FAD
1.1.1.302
RIB2
2,5-Diamino-6-(5-phospho-D-ribitylamino)-pyrimidin-4(3H)-one
3.5.1.102
3.5.4.29
2-Amino-5-formylamino-6-(5-phospho-ribosylamino)-pyrimidin-4(3H)-one
1.5.1.30
1.5.1.36
1.5.1.41
Reduced riboflavin
1.5.1.36
1.5.1.38
1.5.1.39
1.5.1.42
FMNH2
1.5.1.36
1.5.1.37
1.5.1.45
FADH2
1.13.11.79
Terpenoid backbone biosynthesis
Dimethylallyl-PP
3.6.1.76
3.6.1.-
Dimethylallyl-P
2.7.1.-
Prenol
2.5.1.129
Prenylated FMNH2
Dimethyl-benzimidazole
Porphyrin metabolism
Data on KEGG graph
Rendered by Pathview

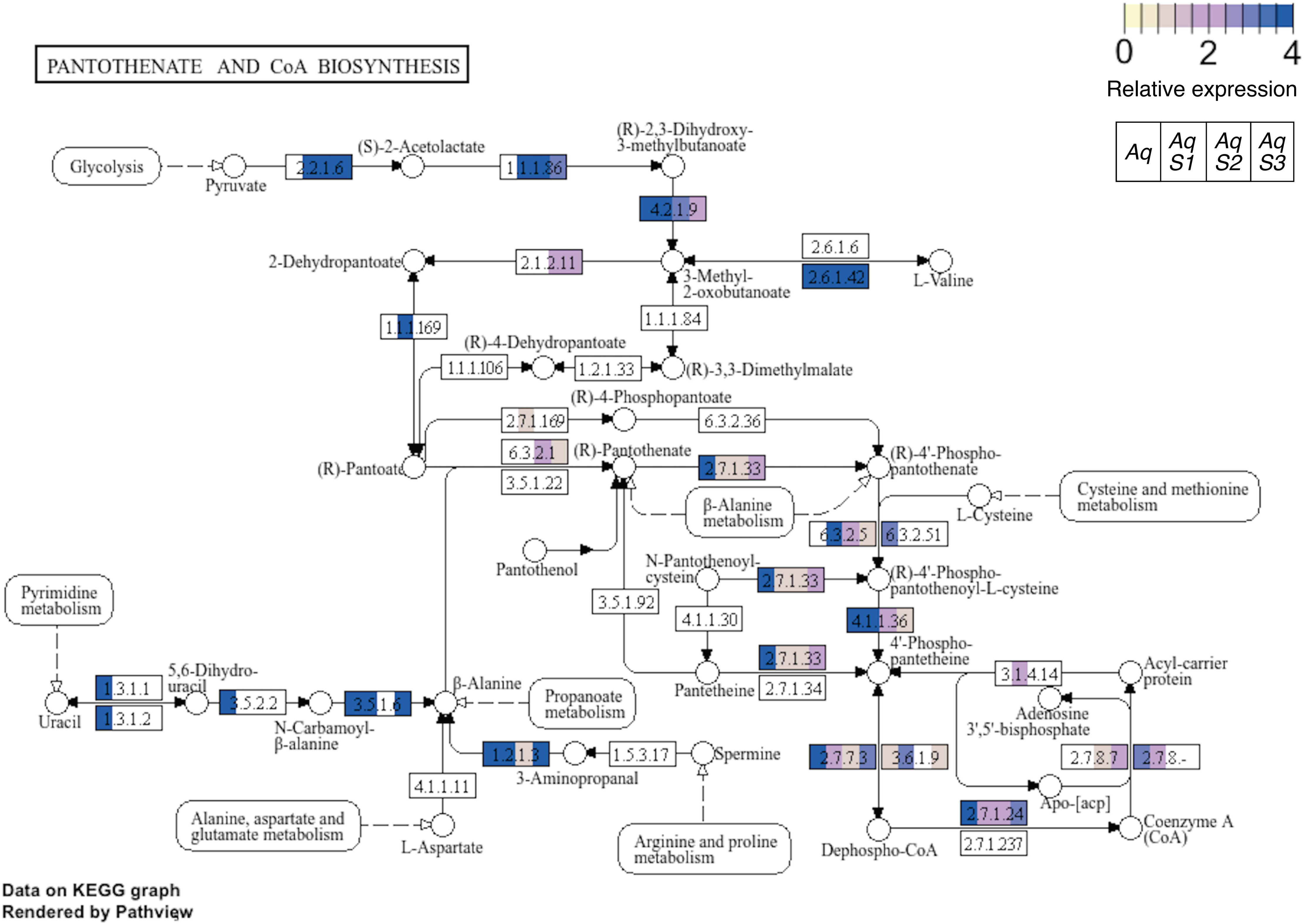
PANTOTHENATE AND CoA BIOSYNTHESIS
0 2 4
Relative expression
Aq
Aq S1
Aq S2
Aq S3
Glycolysis
Pyruvate
2.2.1.6
(S)-2-Acetolactate
1.1.1.86
(R)-2,3-Dihydroxy-3-methylbutanoate
4.2.1.9
2-Dehydropantoate
2.1.2.11
3-Methyl-2-oxobutanoate
2.6.1.6
2.6.1.42
L-Valine
1.1.1.169
1.1.1.84
(R)-4-Dehydropantoate
1.1.1.106
1.2.1.33
(R)-3,3-Dimethylmalate
(R)-4-Phosphopantoate
2.7.1.169
6.3.2.36
6.3.2.1
(R)-Pantothenate
(R)-Pantoate
3.5.1.22
2.7.1.33
(R)-4'-Phospho-pantothenate
Cysteine and methionine metabolism
β-Alanine metabolism
L-Cysteine
6.3.2.5
6.3.2.51
Pantothenol
N-Pantothenoyl-cysteine
2.7.1.33
(R)-4'-Phospho-pantothenoyl-L-cysteine
Pyrimidine metabolism
3.5.1.92
4.1.1.30
4.1.1.36
4'-Phospho-pantetheine
2.7.1.33
Acyl-carrier protein
5,6-Dihydro-uracil
β-Alanine
Pantetheine
2.7.1.34
3.1.4.14
1.3.1.1
3.5.2.2
3.5.1.6
Propanoate metabolism
Adenosine 3',5'-bisphosphate
Uracil
1.3.1.2
N-Carbamoyl-β-alanine
1.2.1.3
1.5.3.17
Spermine
2.7.7.3
3.6.1.9
2.7.8.7
2.7.8.-
3-Aminopropanal
4.1.1.11
Apo-[acp]
2.7.1.24
Alanine, aspartate and glutamate metabolism
L-Aspartate
Arginine and proline metabolism
Dephospho-CoA
2.7.1.237
Coenzyme A (CoA)
Data on KEGG graph
Rendered by Pathview

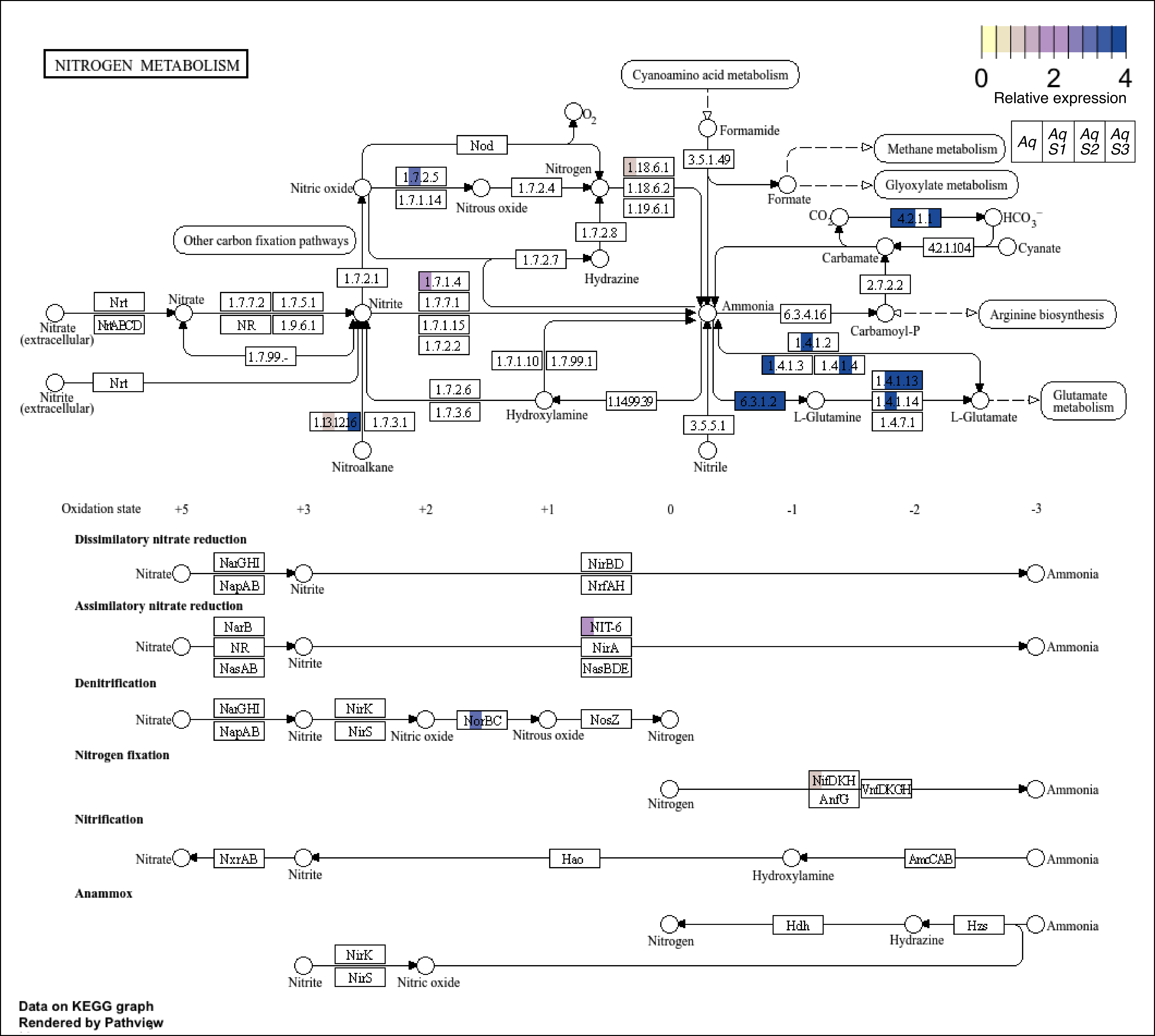
NITROGEN METABOLISM
0
2
4
Relative expression
Aq
Aq S1
Aq S2
Aq S3
Cyanoamino acid metabolism
Formamide
3.5.1.49
Formate
Methane metabolism
Glyoxylate metabolism
Nod
$O_2$
Nitrogen
Nitric oxide
1.7.2.5
1.7.1.14
Nitrous oxide
1.7.2.4
1.18.6.1
1.18.6.2
1.19.6.1
1.7.2.8
1.7.2.7
Hydrazine
Other carbon fixation pathways
$CO_2$
4.2.1.1
$HCO_3^-$
Carbamate
4.2.1.104
Cyanate
2.7.2.2
1.7.2.1
1.7.1.4
1.7.7.1
1.7.1.15
1.7.2.2
Nrt
NrtABCD
Nitrate
Nitrate (extracellular)
1.7.7.2
1.7.5.1
NR
1.9.6.1
Nitrite
1.7.99.-
Ammonia
6.3.4.16
Carbamoyl-P
Arginine biosynthesis
Nitrite (extracellular)
1.4.1.2
1.4.1.3
1.4.1.4
1.7.1.10
1.7.99.1
1.4.1.13
1.4.1.14
1.4.7.1
1.7.2.6
1.7.3.6
1.14.99.39
6.3.1.2
L-Glutamine
L-Glutamate
Glutamate metabolism
Hydroxylamine
1.13.12.16
1.7.3.1
3.5.5.1
Nitroalkane
Nitrile
Oxidation state
+5
+3
+2
+1
0
-1
-2
-3
Dissimilatory nitrate reduction
NarGHI
NapAB
NirBD
NrfAH
Assimilatory nitrate reduction
NarB
NasAB
NIT-6
NirA
NasBDE
Denitrification
NirK
NirS
NorBC
NosZ
Nitrogen fixation
NifDKH
AnfG
VnfDKGH
Nitrification
NxrAB
Hao
AmoCAB
Anammox
Hdh
Hzs
Data on KEGG graph
Rendered by Pathview

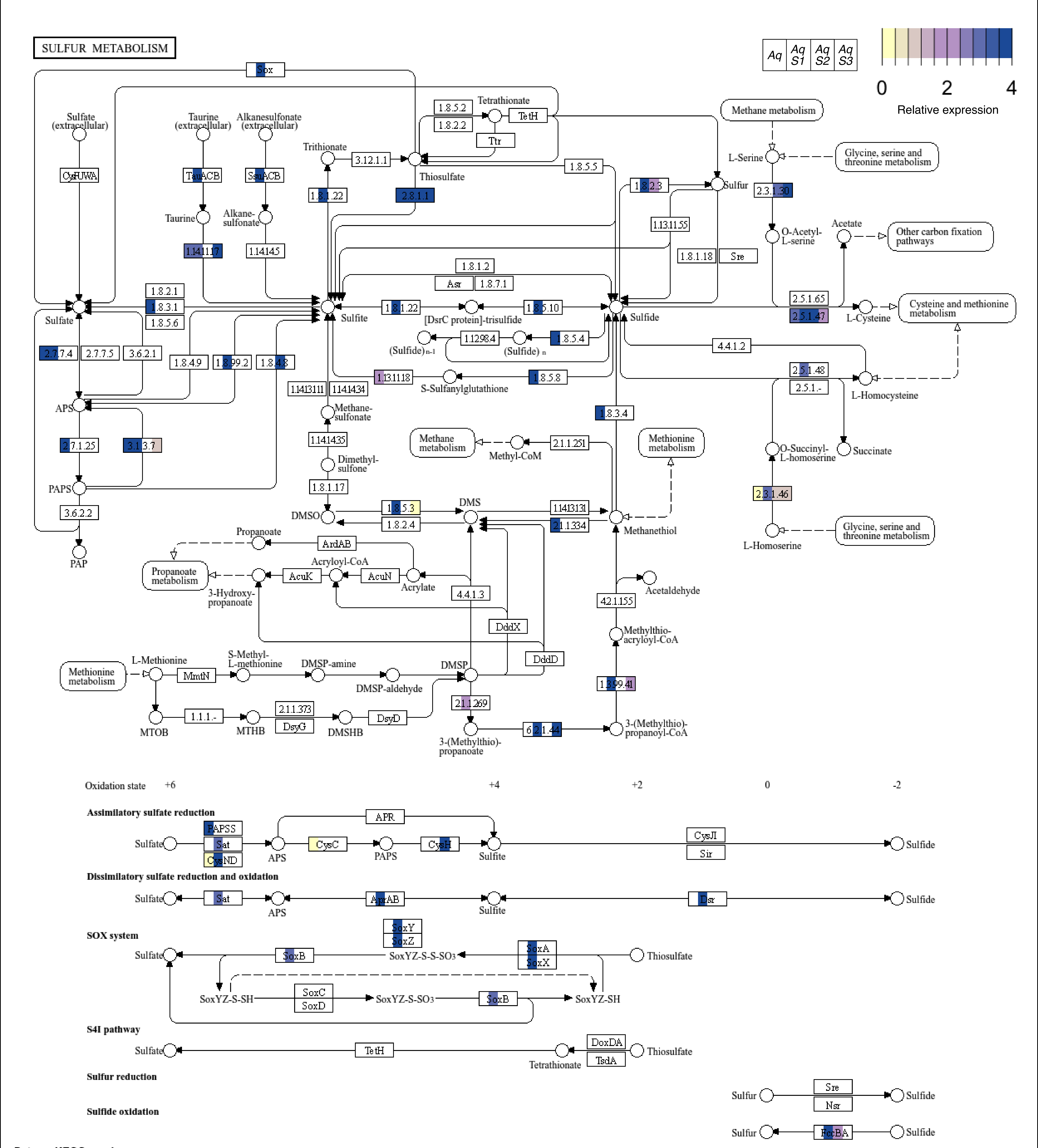


Data on KEGG graph
Rendered by Pathview

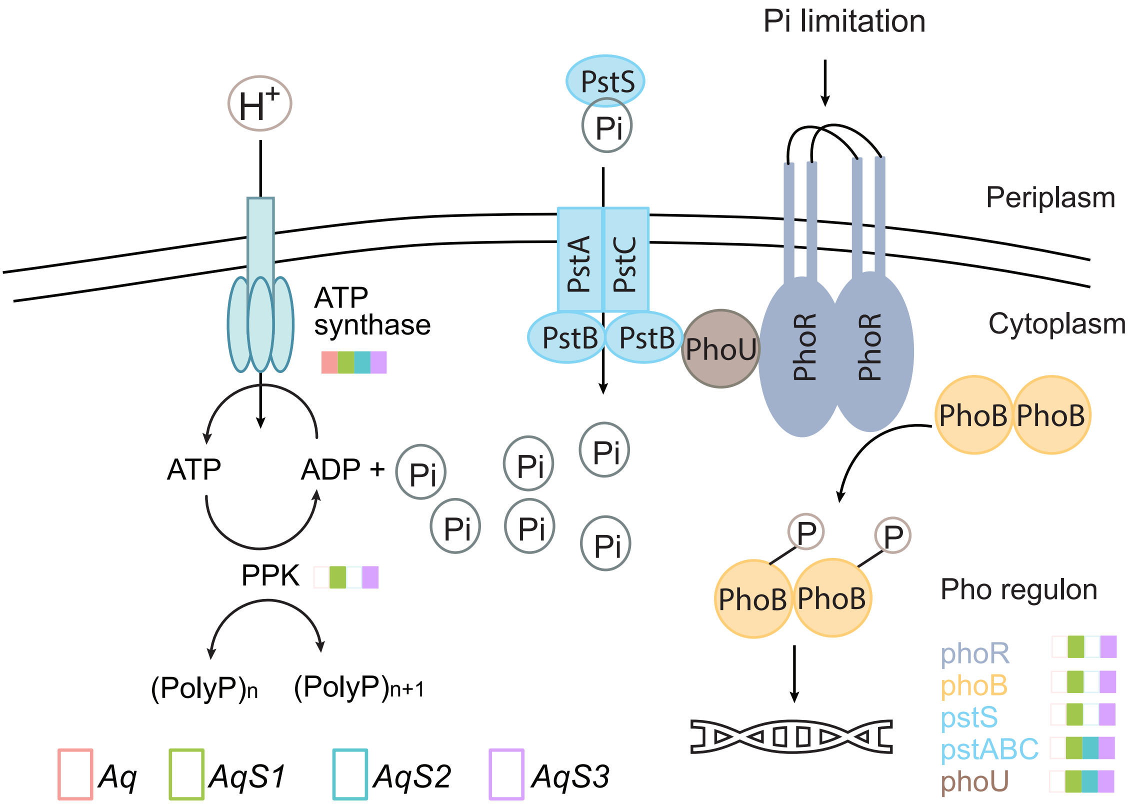

Pi limitation
PstS
Pi
H+
Periplasm
PstA
PstC
PstB
PstB
PhoU
PhoR
PhoR
Cytoplasm
ATP
synthase
PhoB
PhoB
ATP
ADP +
Pi
Pi
Pi
Pi
Pi
Pi
Pi
P
P
PPK
PhoB
PhoB
Pho regulon
phoR
phoB
pstS
pstABC
phoU
(PolyP)n
(PolyP)n+1
Aq
AqS1
AqS2
AqS3
